# Immersive virtual reality interferes with default head-trunk coordination strategies in young children

**DOI:** 10.1101/2020.10.14.338749

**Authors:** Jenifer C. Miehlbradt, Luigi F. Cuturi, Silvia Zanchi, Monica Gori, Silvestro Micera

**Affiliations:** Bertarelli Foundation Chair in Translational Neuroengineering, Center for Neuroprosthetics, École Polytechnique Fédérale de Lausanne, 1202 Geneva, Switzerland; Brain Electrophysiology Attention Movement Laboratory, Institute of Psychology, University of Lausanne, 1015 Lausanne, Switzerland; Unit for Visually Impaired People, Center for Human Technologies, Fondazione Istituto Italiano di Tecnologia, 16163 Genova, Italy; DIBRIS Department, Università di Genova, 16163 Genova, Italy; The BioRobotics Institute, Scuola Superiore Sant’Anna, 56025 Pontedera, Italy

## Abstract

The acquisition of postural control is an elaborate process, which relies on the balanced integration of multisensory inputs. Current models suggest that young children rely on an ‘en-block’ control of their upper body before sequentially acquiring a segmental control around the age of 7, and that they resort to the former strategy under challenging conditions. While recent works suggest that a virtual sensory environment alters visuomotor integration in healthy adults, little is known about the effects on younger individuals.

Here we show that this coordination pattern is disrupted by an immersive virtual reality framework where a steering role is assigned to the trunk, which causes 6- to 8-year-olds to employ an ill-adapted segmental strategy. These results provide an alternate trajectory of motor development and emphasize the immaturity of postural control at these ages.

## Introduction

Coordinated motor behavior and efficient integration of stimuli from different sensory modalities are necessary for successful interactions with the surrounding environment (*1*). The development of these abilities follows a long-lasting and elaborate process, starting long before birth and extending into early adulthood. At the motor development level, the skills are usually grouped into two categories. First, gross motor skills comprise postural control and locomotion and require the use of axial and proximal muscles. The maturation of these abilities shows a steep increase until the age of 2 years and continues to refine until later childhood (*2–5*). Conversely, fine motor skills include precise actions such as functional hand movements, but also require multisensory integration such as hand-eye coordination. The time course of fine motor development typically extends over a more extended time period and adult patterns are generally not observed before late childhood (*6, 7*)

The acquisition of a steady posture is a prerequisite for goal-directed behaviors such as reaching from a sitting position or locomotion (*1, 6*). According to the ontogenetic model of postural development during childhood described by Assaiante et al., two main principles guide the selection of a given balance strategy: the choice of a stable reference, which shifts from the pelvis to the head (*1, 8*), and the gradual mastery of the involved degrees of freedom (DOF) (*1, 9, 10*). The coordination strategy evolves from an ‘en-block’ behavior, which minimizes the number of DOF to be controlled (*11, 12*) to a fully articulated strategy, where each DOF is controlled individually. Mature, multi-jointed patterns are acquired at different ages, depending on the involved joint and task characteristics. During locomotion, the ‘en-block’ stabilization has been observed from the acquisition of an upright stance until 6 years, while children aged 7 and older started to display a segmental control (*10*). Similarly, rigid forearm-trunk coupling was observed until 6 years both during voluntary trunk movements and in response to trunk perturbations (*13*). Instead, in a reaching task, adult head-trunk-arm coordination patterns were observed in children as young as 2-3 years old for movements in the pitch plane and from 4 years onwards in the roll and yaw planes (*14*). Yet, the activity and temporal recruitment of postural muscles appear to reach mature levels only after the age of 11 (*8*). The ability to decouple head and trunk movements proves to be particularly useful when having to avoid or circumvent an obstacle while walking, where anticipatory head movements were observed from 5.5 years onwards, while younger children displayed a rigid head-trunk connection (*15*). Children thus first build a repertoire of postural strategies, before learning how and when to adequately implement them.

Nevertheless, successful postural stabilization does not only involve appropriate multi-jointed coordination but also requires the integration of the information provided by different sensory modalities. The Bayesian model of multisensory integration suggests that adults fuse redundant sensory inputs in a statistically optimal way by weighting the sources according to their uncertainty (*16, 17*). The ability to combine different cues to obtain more precise estimates of one’s surroundings appears late in childhood development (*18, 19*), that is, after the individual modalities have matured (*20, 21*), unless additional feedback on the reliability of each cue is provided (*22*). Younger children will thus favor the information provided by the modality with the highest context-dependent reliability (*19, 23*). In the case of postural control, children and adolescents until 15 years standing on an oscillating platform displayed better stabilization with open than with closed eyes, thus indicating a strong reliance on vision (*3, 24*). The display of optic flow patterns to elicit automatic postural movements led to stronger responses in children and adolescents when compared to adults, and the ability to stabilize these movements improved with age until late adolescence (*25*). This effect was further enhanced when the participants were standing on a sway-referenced platform (*26, 27*). When standing on the unstable platform, which attenuates the proprioceptive feedback, adults use primarily vestibular information to stabilize their posture, and this ability matures only during late adolescence (*26*).

Interestingly, children aged 7–10 years have been shown to display spatiotemporal muscle activation patterns similar to those observed in adults in response to platform oscillations (*28*), revealing an earlier development of automatic postural responses. Similarly, the predominance of visual cues over self-motion has been observed in children up to 11 years in a navigation task (*29, 30*). The late maturation of visual-vestibular and visual-proprioceptive integration has been correlated with the individual development of these modalities when these are presented in conflict.

The reliance on visual cues can be further challenged by the use of immersive VR, where the participants are immersed in a digital environment through a head-mounted display (HMD). This paradigm led to stronger sensory recalibration (*31*) and recruited different adaptation mechanisms (*32*) than non-immersive sensory alterations. Thanks to the recent development of lightweight HMDs, the use of VR has expanded to numerous applications designed for children, including neurodevelopmental research (*30, 33–35*), neurorehabilitation (*36–39*), or distraction from painful medical procedures (*40, 41*). Yet, the majority of these applications offer none or limited interactions with the virtual environment. Therefore, with the exception of two studies showing that children displayed stronger and longer-lasting responses than teenagers to prism adaptation in immersive VR (*42*), but generally tolerate this kind of environment (*43*), little is known about how children integrate the visual information of the simulated world.

We previously developed a body-machine interface for the immersive control of a first-person view (FPV) flight simulator relying on simple trunk movements, which was rapidly mastered by healthy adults (*44*). Here, we first evaluated the ability of school-aged children to control this interface using either their head or their torso, and we assessed the intersegmental coordination patterns which emerged during the execution of this task (Study 1). To further investigate the underlying behaviors, we assessed the development of the head and torso proprioception during a virtual joint angle reproduction task (JAR, Study 2).

## Results

### Study 1

In the first study, the participants were equipped with a HMD through which they were immersed in a virtual scenario representing a flight on a bird’s back along a path represented by a series of coins to catch (Fig.1A). The trajectory of the flight simulator was controlled either by head movements or torso movements. Continuous tracking of the head movements also enabled a dynamic adaptation of the field of view, allowing the users to look around in the virtual environment (Fig. 1B). Steering with torso movements, therefore, required decoupling of vision and steering commands, whereas these aspects were tied in the head-controlled trials.

**Fig. 1:**
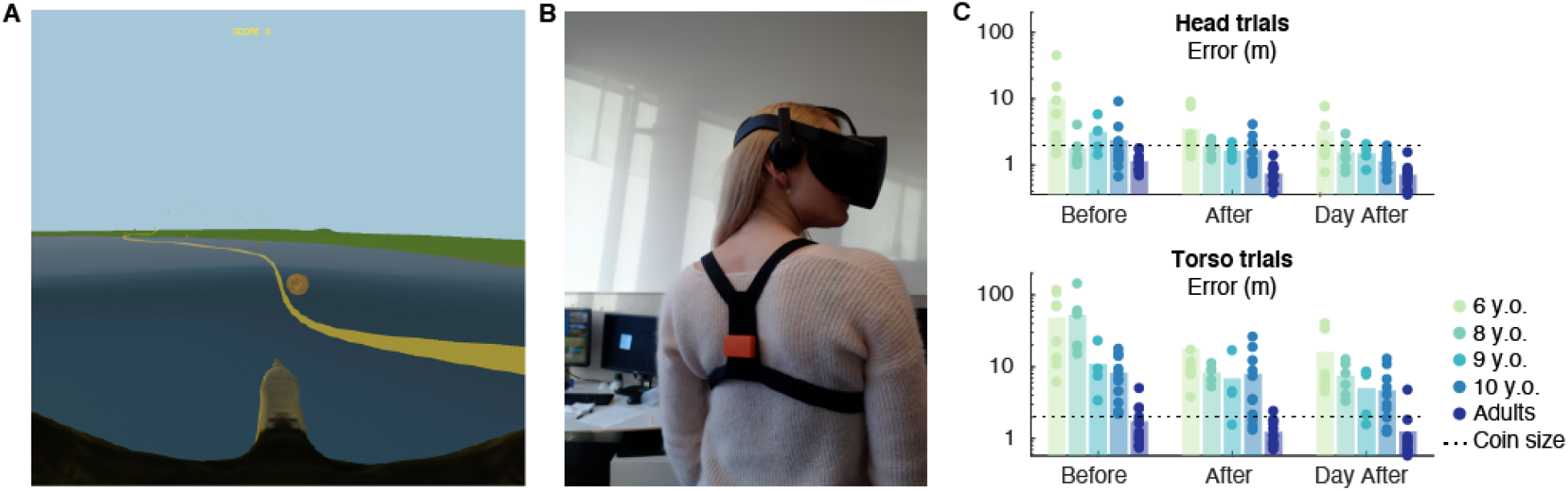
Experimental setup and task performance. A Virtual environment, as seen by the participant, representing the coins to catch and an underlining ideal trajectory depicted by the yellow line. B Experimental apparatus worn by the participants, consisting of a HMD and an IMU held in place in the back by a harness. C Performance on the navigation task, computed as the average distance to the coin centre (error). Dots represent the average error for each individual participant, bars the average across participants. N = 9 (6 y.o.), 8 (8 y.o.), 4 (9 y.o.), 11 (10 y.o.), 13 (adults). See Tables S3 and S4 for details of the statistical analyses.

#### Controlling body part and age affect steering performance

We assessed the steering performance as the average distance to the center of the coins (*44, 45*) during three phases: before and after training (*Before* and *After,* see Methods), and on the subsequent day (*Day After*). A repeated measures ANOVA revealed a significant effect of Age (F(4,35) = 7.45, p < 0.001, η_p_^2^ = 0.460), Control (F(1,35) = 29.52, p < 0.001, η_p_^2^ = 0.457) and Phase (F(2,70) = 15.44, p < 0.001, η_p_^2^ = 0.306), as well as significant Age:Phase (F(8,70) = 4.41, p = 0.003, η_p_^2^ = 0.335), Age:Control (F(4,35) = 5.97, p < 0.001, η_p_^2^ = 0.405) and Age:Phase:Control (F(8,70) = 4.21, p = 0.003, η_p_^2^ = 0.325) interactions (Fig. 1C).

Post-hoc Tukey tests revealed that 6-year-olds performed better in the head-than in the torso-controlled trials in all phases (*Before*: p = 0.002, *d* = 1.17; *After*: p = 0.009, *d* = 0.83; *Day After*: p < 0.001, *d* = 1.17). This difference was also significant for 8-year-olds *Before* (p < 0.001, *d* = 1.45), but not during the other phases.

When steering with their torso, 6-year-olds performed better *After* than *Before* training (p = 0.013, *d* = 0.85) and on the *Day After* than *Before* (p = 0.014, d = 0.97). The same improvement was observed in 8-year-olds between the evaluations *Before* and *After* training (p = 0.001, *d* = 1.26) and from *Day After* compared to *Before* training (p = 0.002, *d* = 1.28). Interestingly, large effect sizes suggest that 9-year-olds and adults improved their steering precision in head-controlled trials from *Before* to *After* and *Day After* (Table S1).

In the torso-controlled trials, 6-year-olds showed significantly lower performance than 10-year-olds *Before* training (p = 0.023, *d* = 1.34) and on *Day After* (p = 0.02, *d* = 1.07) and than adults in all phases (*Before*: p = 0.006, *d* = 1.66; *After*: p = 0.042, *d* = 1.09; *Day After*: p = 0.001, *d* = 1.1.57). Likewise, 8-year-olds performed worse than 10-years-olds and the adults *Before* training (p = 0.015, *d* = 1.45 and p = 0.005, *d* = 1.78 respectively). In the head-controlled trials, 6-year-olds displayed higher errors than the adults *After* training (p = 0.001, *d* = 1.55), and than 10-year-olds and the adults on *Day After* (p = 0.013, *d* = 1.07 and p = 0.002, *d* = 1.52 respectively). Non-significant differences with large effect sizes suggest a gradual development of head-torso motor patterns, particularly between the two older children groups and adults (Table S2).

#### Segmental coordination and torso involvement differ between the torso and head trials

Principal Component Analysis (PCA) applied to all the recorded trials revealed that the first principal component (PC) accounted for 34% of the dataset’s variability and separated the head-from the torso-controlled trials (p < 0.001, Fig. 2A). The kinematic variables displaying normalized loadings > 0.75 represented torso movements (Cluster 1) and head-torso coordination (Cluster 2, see Fig. 2B).

**Fig. 2:**
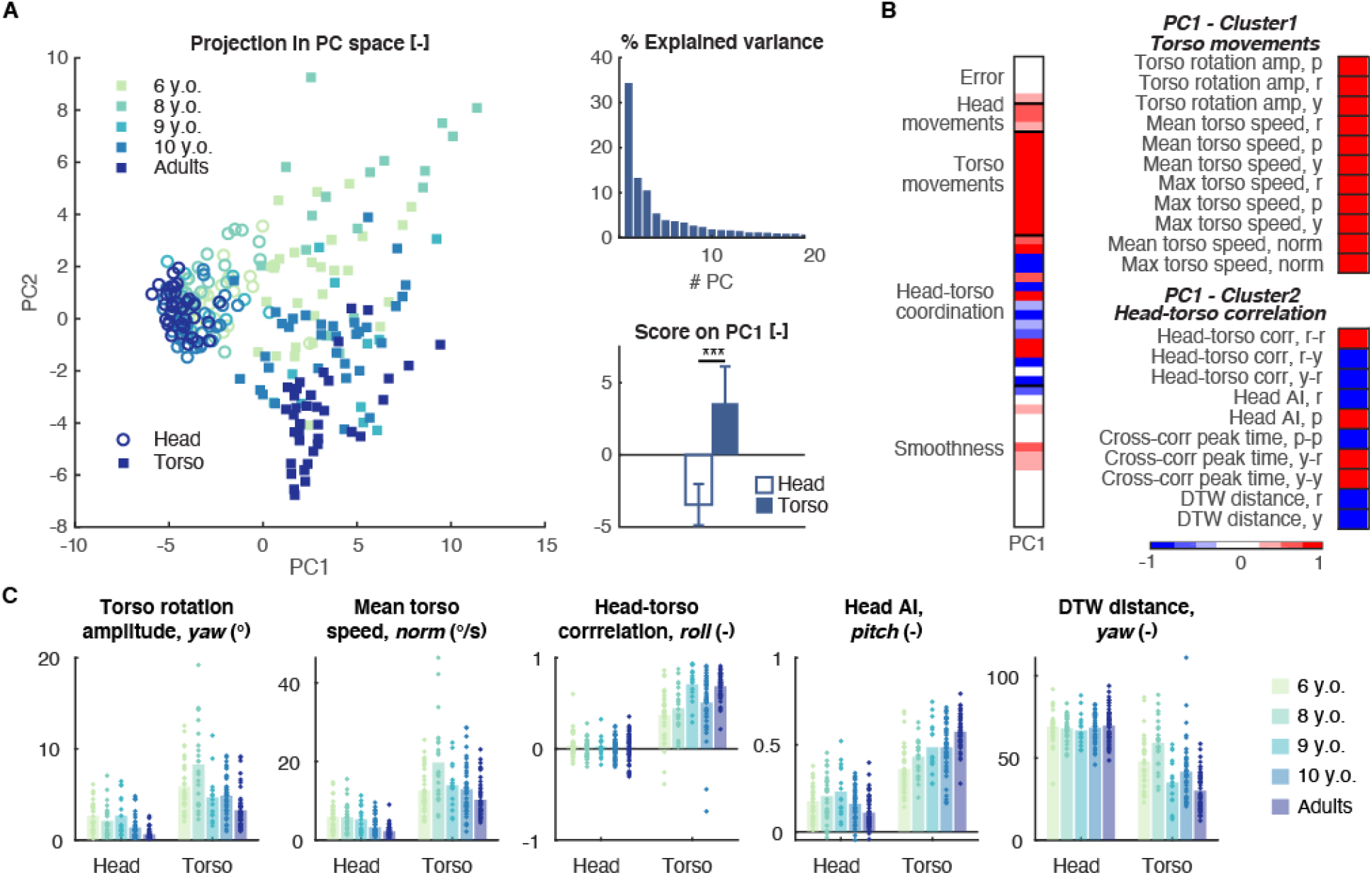
Segmental coordination and torso involvement differ between torso and head trials. A PCA applied to the data collected on all trials. The projection of the data in the space spanned by the first two PCs displays a control-based separation along the first component (left) representing 34% of the overall variance (top right). This division was confirmed by a t-test (bottom left, mean + SEM). B Normalized loadings of the descriptive variables on the first PC (left) and variables with absolute loadings higher than a threshold of 0.75 grouped into functional clusters. C Representative variables selected from the functional clusters with significant effect of Control (see also Table S4). B: Before, A: After, DA: Day After

Repeated measures ANOVAs revealed a significant effect of Control on all identified variables (Table S3). In particular, torso movements were executed with larger yaw amplitude (p < 0.001, η_p_^2^ = 0.67) and higher average velocity (p < 0.001, η_p_^2^ = 0.79) in the torso-controlled trials (Fig. 2C). Head movements were more similar to trunk movements in torso-than in head-controlled trials, as assessed by the head-torso correlation in the roll plane (p < 0.001, η_p_^2^ = 0.89) or the dynamic time warp (DTW) distance between both segments in the yaw plane (p < 0.001, η_p_^2^ = 0.82). Interestingly, the higher pitch head anchoring index (AI) in the torso-controlled trials (p < 0.001, η_p_^2^ = 0.85) reveals that the head is preferentially stabilized to the external space than to the trunk in these trials.

#### Efficient selection of head-torso coordination strategy develops with age

To extract the specific variability inherent to torso steering, we repeated the procedure described above, using only the data from the corresponding trials. On this partial dataset, PCA revealed an age-based separation in the space spanned by the first two PCs, accounting respectively for 25.91% and 19.38% of the total variance (Fig. 3A). Individually, both PC1 and PC2 showed a decreasing trend with age (Fig. 3A).

**Fig. 3:**
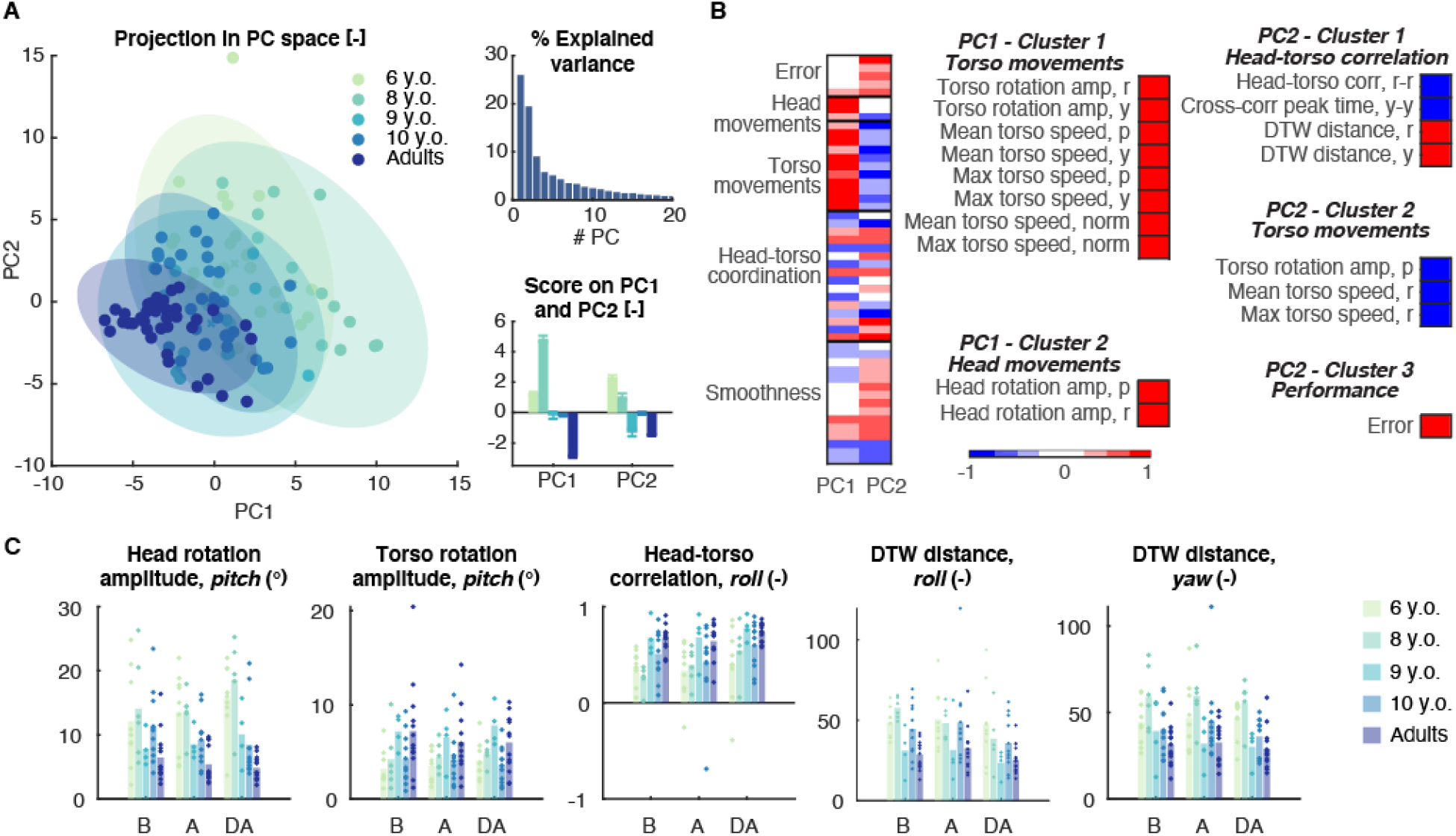
Efficient selection of head-torso coordination strategy develops with age. A PCA applied to the data collected on torso-controlled trials. The projection of the data in the space spanned by the first two PCs displays an age-based separation along the first two components (left) representing respectively 26% and 19% of the overall variance (top right). Group means of the scores on the first two PCs (bottom left, mean + SEM). B Normalized loadings of the descriptive variables on the first PC (left) and variables with absolute loadings higher than a threshold of 0.75 grouped into functional clusters. C Representative variables selected from the functional clusters with significant effect of Age (see also Table S5). B: Before, A: After, DA: Day After

The selection of relevant descriptive variables yielded five functional clusters: Cluster 1 (PC1) and Cluster 2 (PC2) holding variables describing the torso movements, Cluster 2 (PC1) corresponding to head movements, Cluster 1 (PC1) characterizing head-torso correlation and finally Cluster 3 (PC2) containing only the error (Fig. 3B). All the identified variables showed a significant effect of Age and/or Age:Phase interaction (Table S4). Younger children displayed larger vertical head movements (p = 0.004, η_p_^2^ = 0.42, Fig. 3C) and smaller torso movements (p = 0.003, η_p_^2^ = 0.44). Remarkably, the similarity between head and torso movements augmented with age, as revealed by the increased correlation in the roll plane (p = 0.01, η_p_^2^ = 0.43) or the DTW distance in the roll (p = 0.005, η_p_^2^ = 0.39) and yaw planes (p = 0.003, η_p_^2^ = 0.44).

#### Torso involvement in head-controlled trials decreases with age

For head-controlled trials, PCA revealed a soft age-based separation along with the first principal component, accounting for 25% of the total variance (Fig. 4A). Clustering the variables with normalized loadings larger than 0.75 yielded one single cluster describing torso movements (Fig. 4B). All the identified variables showed a significant effect of Age and/or Age:Phase interaction (Table S5). The amplitude of the torso movements decreased with age in the pitch (p = 0.016, η_p_^2^ = 0.31, Fig. 4C) and yaw planes (p = 0.015, η_p_^2^ = 0.32), as well as the average (p = 0.016, η_p_^2^ = 0.3) and maximal torso velocity (p = 0.015, η_p_^2^ = 0.32).

**Fig. 4:**
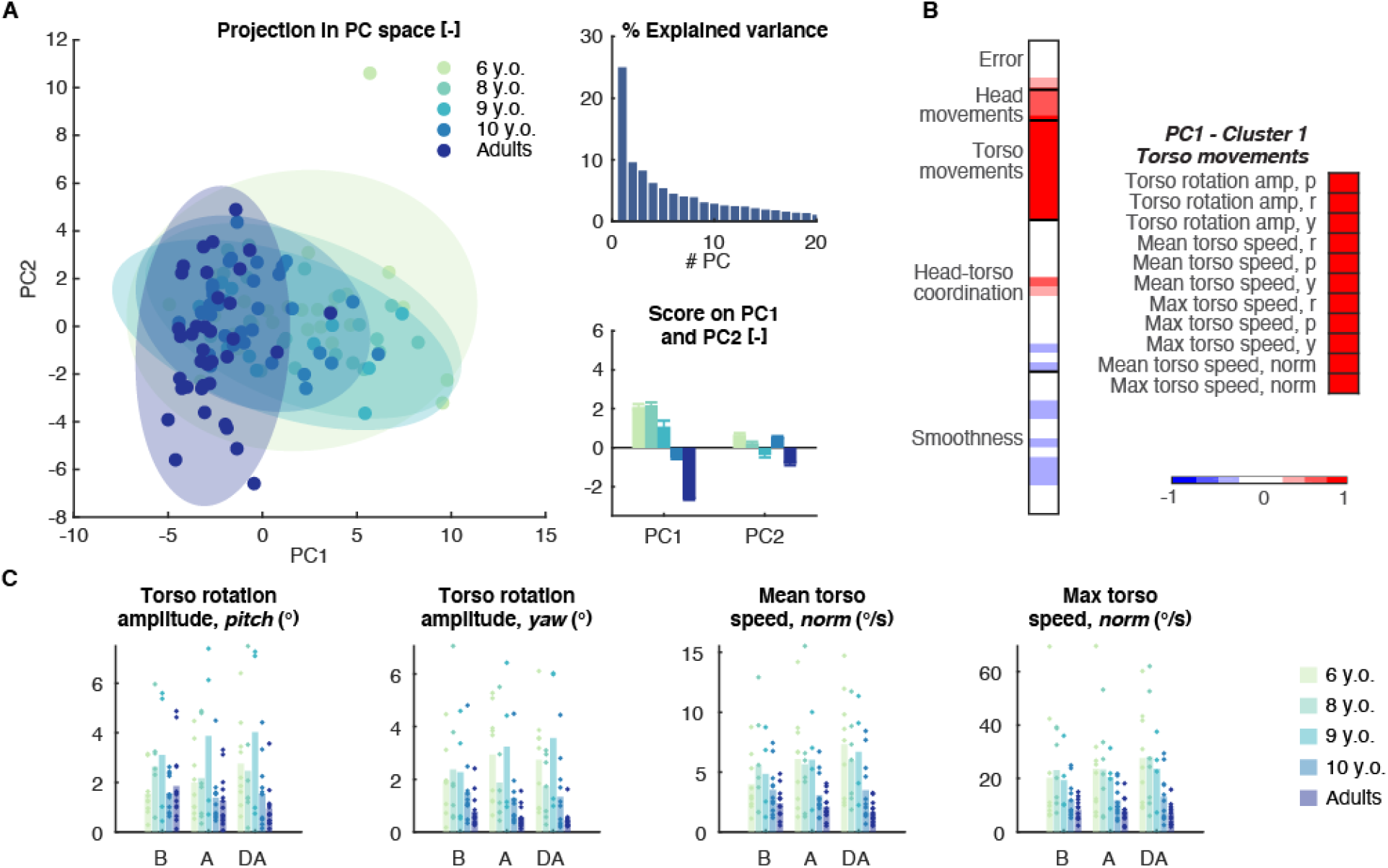
Torso involvement in head-controlled trials decreases with age. A PCA applied to the data collected on head-controlled trials. The projection of the data in the space spanned by the first two PCs displays an age-based separation along the first component (left) representing 25% of the overall variance (top right). Group means of the scores on the first two PCs (bottom left, mean + SEM). B Normalized loadings of the descriptive variables on the first PC (left) and variables with absolute loadings higher than a threshold of 0.75 grouped into functional clusters. C Representative variables selected from the functional clusters with significant effect of Age (see also Table S6). B: Before, A: After, DA: Day After

### Study 2

To further elucidate the mechanisms underlying the observed behavior, in particular the importance of mature and reliable proprioceptive inputs under altered visual feedback, we designed a second study in which the participants were immersed in a virtual landscape as previously and asked to execute a JAR test using their head or their torso. The JAR paradigm is an active test for proprioception that reflects the functional use of this sensory pathway and relies on kinaesthetic memory (*46, 47*), a necessary competence for the proficient use of the flight simulator tested in Study 1.

#### Error

We first evaluated the angle reproduction error under three conditions: *Feedback*, where a line indicated the current angle of the tested body part, *Still,* where the feedback line was removed, and *Forward*, where a constant forward speed was simulated and no feedback line was displayed. A repeated-measures ANOVA showed a significant effect of Age (F(3,35) =7.99, p < 0.001, η_p_^2^ = 0.406) and Control (F(1,35) = 21.19, p < 0.001, η_p_^2^ = 0.377), and significant Age:Control (F(3,35) = 5.24, p = 0.004, η_p_^2^ = 0.446) and Age:Control:Condition interactions (F(3,35) = 3.99, p = 0.003 η_p_^2^ = 0.255). A posthoc analysis revealed that in the absence of feedback, all age groups except the 6-year-olds increased their error when using their torso compared to the head trials, overestimating their position in the former case and underestimating it in the latter (see tables S6 and S7 for details). This was particularly the case in the *Forward* condition for the 8- and 10- year-olds and the adults, and in the *Still* condition for the 10-year-olds and the adults (Fig. 5A). The youngest participants in turn failed to reach the target angle with both body parts.

**Fig. 5:**
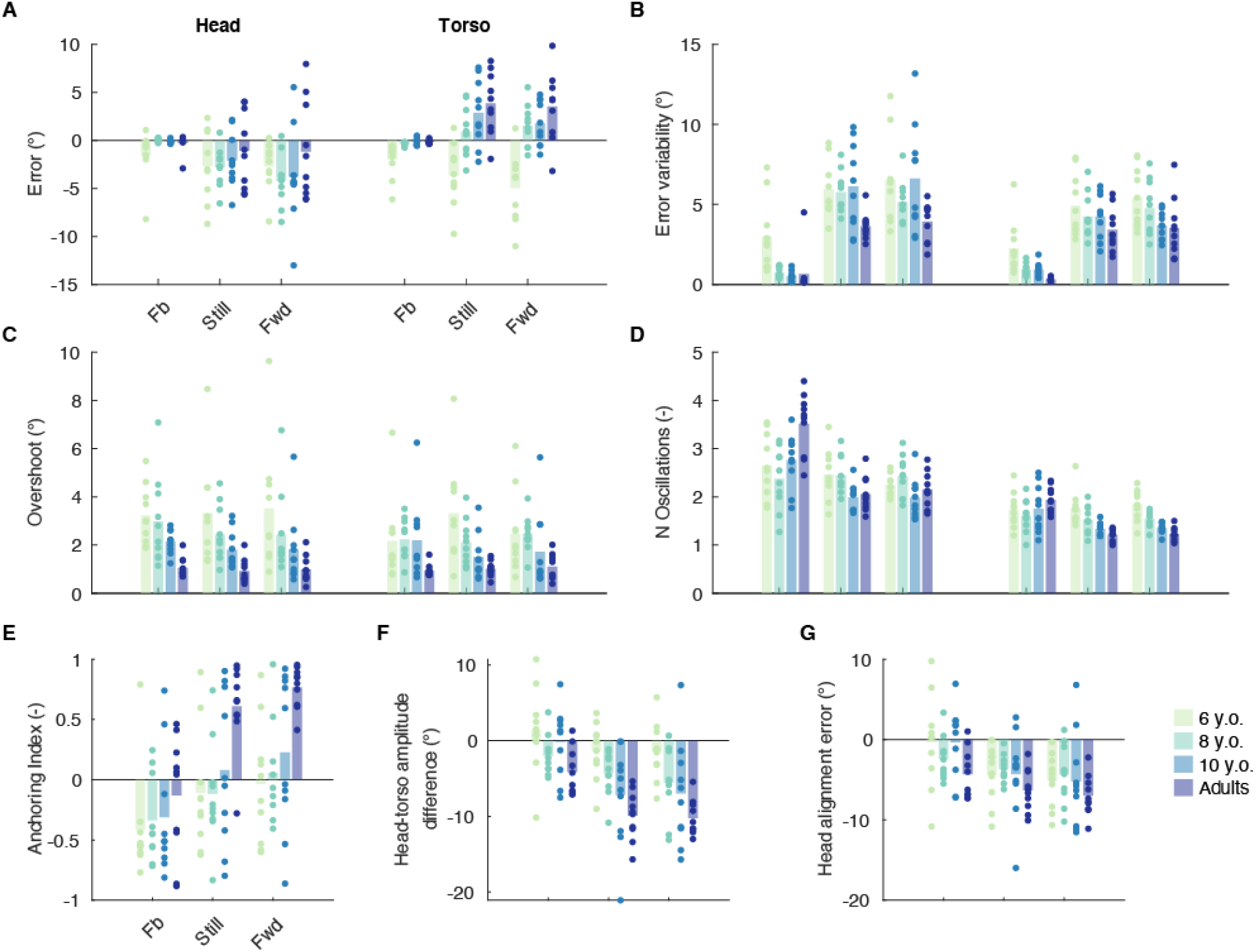
Joint angle reproduction (JAR) test. A-D Head and torso trials, E-G Torso trials only. A Signed error at final orientation, positive values indicate final positions exceeding the target angle. B Variability (standard deviation) of the final error. C Overshoot. D Number of oscillations around the final position. E Head anchoring index (AI), with AI = 1 meaning complete independence of head and torso. F Difference of head and torso final orientation, negative values indicate that the head orientation is smaller than the torso orientation. G Difference between final head orientation and target orientation. Dots represent the average error for each individual participant, bars the average across participants. N = 10 for each age group. See Tables S6 – S9 for details of the statistical analyses. FB: Feedback, Fwd: Forward, see text for description of the conditions.

**Fig. 6:**
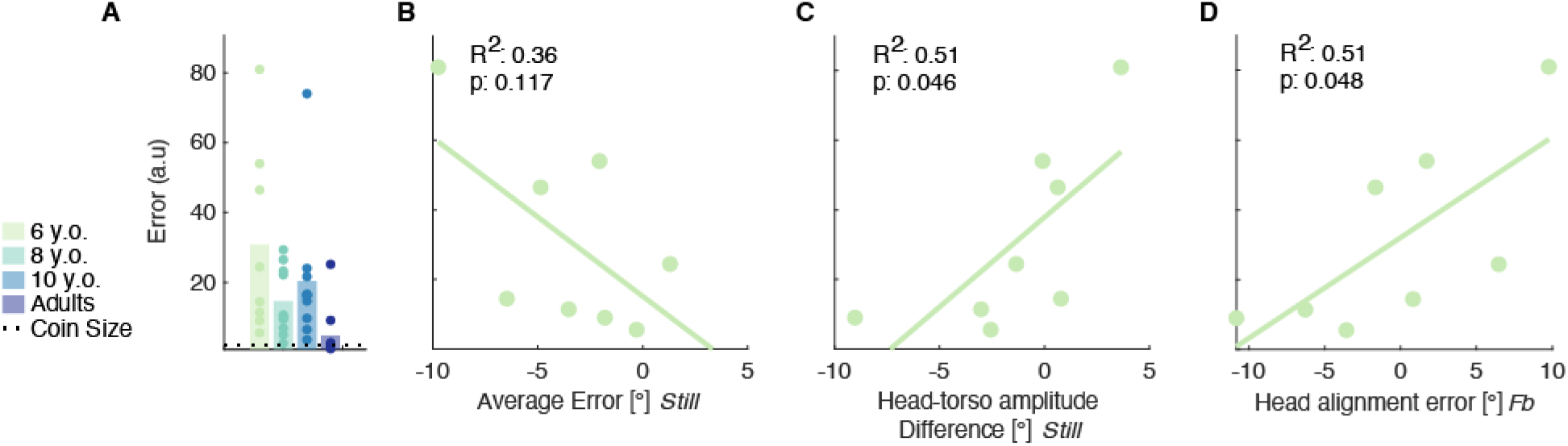
Prediction of simulator steering performance from JAR test. A Simulator steering performance during a unique torso-controlled session. B-D Regression analysis performed using the data of the 6-year-olds, recorded during torso trials of the JAR test. B Signed error at final orientation (see Figure 5A). C Difference between final head orientation and target orientation (see Figure 5F). D Difference between final head orientation and target orientation (see Figure 5G). Dots represent the average error for each individual participant, bars the average across participants. N = 8 (6 y.o.), 10 (8 y.o.), 10 (10 y.o.), 10 (adults). FB: Feedback, see text for description of the conditions.

We found a significant effect of Age (F(3,35) = 9.41, p < 0.001, η_p_^2^ = 0.446), Condition (F(2,70) = 152.40, p < 0.001, η_p_^2^ = 0.813) and Control (F(1,35) = 10.98, p = 0.002, η_p_^2^ = 0.0.239) on the variability of the error, but none of the interactions involving Age were significant. Posthoc tests revealed that adults showed significantly less variability than 6- and 10-year-olds (p < 0.001, *d* = 0.92 and p = 0.028, *d*= 0.45 respectively, Fig. 5B).

#### Movement strategy

We next evaluated the selected movement strategy through the overshoot with respect to the final position and the number of oscillations around this angle. We found a significant effect of Age (F(3,35) = 5.53 p =0.003, η_p_^2^ = 0.322) and Control (F(1,35) = 4.55 p =0.04, η_p_^2^ = 0.115) on the overshoot, with 6- and 8-year-olds exceeding their final position by a larger extent than adults (p = 0.03, *d* = 1.40 and p = 0.031, *d* = 1.53 respectively, Fig. 5C).

Assessing the number of oscillations around the final position, we found a significant effect of Condition (F(2,35) = 36.07 p < 0.001, η_p_^2^ = 0.508), and Control (F(1,35) = 342.80, p < 0.001, η_p_^2^ = 0.907), and Age:Condition (F(6,70) = 11.97, p < 0.001, η_p_^2^ = 0.506), Age:Control (F(3,35) = 9.41, p = 0.015, η_p_^2^ = 0.0.255), and Age:Condition:Control (F(6,70) = 3.37, p = 0.006, η_p_^2^ = 0.224) interactions. Specifically, we found that adults oscillate more than all children groups when using their head and *Feedback* is provided while 6- and 8-year-olds oscillate more than older children and adults when using their torso in the *Still* and *Forward* conditions. (Fig. 5D, see tables S8 and S9 for details)

#### Head-torso coordination during torso trials

We observed a significant effect of Age, Condition and Age:Condition interaction on the head anchoring index (AI). The 6- and 8-year-olds’ AI was significantly lower than the adult’s in the *Still* (p = 0.013, *d* = −1.67 and p = 0.019, *d* =−1.85 respectively) and *Forward* conditions (p = 0.003, *d* = −2.15 and p = 0.012, *d* =−2.14 respectively, Fig. 5E).

The angular difference between the head and torso orientations at the final position showed a significant effect of Age (F(3,36) = 7.29 p = 0.001, η_p_^2^ = 0.378) and Condition (F(2,72) = 36.07 p < 0.001, η_p_^2^ = 0.508). This difference was significantly smaller for 6-year-olds than adults (p < 0.001, *d* = 1.78, Fig. 5F). Finally, there was only an effect of Condition on the alignment error of the head with the target orientation (F(2,72) = 18.11 p < 0.001, η_p_^2^ = 0.335, Fig. 5G).

#### Effect of optical flow

Interestingly, the effect of the constant optic flow implemented to generate the *Forward* condition when compared with the *Still* condition was significant only for the head AI, regardless of the age group (p = 0.004, *d* = −0.26).

#### Prediction of performance in the flight game

Eventually, we evaluated the relationship between the metrics computed during the JAR test and the performance during one torso-controlled session on the flight simulator. For the 6-year-olds, we found a significant relationship of this performance with the head-torso amplitude difference in the absence of visual feedback (*Still*: R^2^ = 0.51, p = 0.046 *Forward*: R^2^ = 0.60 p = 0.024), as well as with the head alignment error with *Feedback* (R^2^ = 0.51, p = 0.048). None of the regressions were significant for the other age groups. Interestingly, we found no significant relationship between the torso JAR error and the flight performance (*Still:* R^2^ = 0.36, p = 0.117, *Forward:* R^2^ = 0.35, p = 0.120 for the 6-year-olds).

## Discussion

We investigated the development of head-torso coordination when challenged by an alteration of the visual feedback through immersive VR. We first evaluated the ability of children aged 6-10 years and young adults to steer an immersive flight simulator using either their head or their torso (Study 1), followed by a virtual JAR task to further address the behaviors observed during the steering task (Study 2).

All the participants were able to control the flight path using their head in Study 1. 6-year-olds showed lower performances than the oldest children and adults even after practicing the task, but the scores were in a comparable range. When using their torso, 6 and 8-year-olds initially struggled to control the simulator but substantially improved their performance with training. Yet, their average error remained higher than the 10-year-olds’ and adults’. The age-dependent kinematic changes in the head-controlled trials were predominantly due to a decrease of superfluous torso movements. Conversely, the age-related differences in the torso-controlled trials were explained by an increase of the torso movement amplitude, a decrease of the head movements and an increase in the head-torso correlation.

The virtual JAR test carried out in Study 2 revealed that, in the absence of explicit visual feedback, participants aged 8 and older failed to reach the target angle with their head while exceeding it when performing the task with their torso. The younger children instead failed to reach the desired orientation with both body parts, overestimating their displacement in either case. We also observed that in the torso JAR test, older children and adults decoupled their heads from their torso, maintaining the head close to the vertical during sideward trials. Instead, when explicit feedback was given on the torso position, the 6-year-olds had the tendency to overshoot the target orientation with their head. Lastly, we found that for this age group, the amplitude of unnecessary head movements during the torso JAR correlated with their performance in the torso-controlled flight game, while no such relation was found for the amplitude or precision of torso movements.

The comparable performances observed for all age groups in the head-controlled JAR and steering task indicate that children as young as 6 years are able to use and interact with an immersive body-machine interface both for simple and more complex tasks, in line with a recent study (*43*). The earlier maturation of the head control is not surprising, as this condition does not require the mastery of an articulated control of the head-trunk unit, which develops from 7 years onwards (*10*). However, even in this simpler experimental condition, younger children still display a higher error variability and a larger overshoot, confirming the incomplete development of robust internal models as observed in standard experimental frameworks (*2, 48, 49*)

Kinematic analyses of the head-controlled trials showed that the major age-related difference could be attributed to differences in the torso movements, with rotation amplitudes and mean and maximum rotation velocities are decreasing with age. The ability to decouple head from torso movements thus develops along with childhood, confirming previous results obtained during obstacle avoidance during locomotion (*1, 15*), where adults display anticipatory head movements (*15*). However, mature coordination patterns appear later with our experimental setup when compared to simple locomotion. This is in line with observations revealing that developing children tend to increase their head-body stiffness with increasing task difficulty (*9*), and to involve their trunk in situations where such movements are not necessarily required (*50, 51*). In our case, the increased difficulty can be imputed to the use of immersive VR, which provides altered visual information and requires higher cognitive processing abilities to appropriately interpret the displayed environment (*52, 53*).

When the control of the flight game was based on torso movements instead, younger children struggled to use the system, even after practicing the task. Assessing the kinematics during this task and the JAR reveals an underlying twofold behavior. First, the age-related increase of the torso amplitude in the steering task and the evolution of the torso JAR error indicate that the immaturity of the torso proprioception leads younger children to overestimate their torso movements. This complements a previous study showing an increase in torso positioning accuracy with age (*2*). Second, the larger head movements displayed by the younger participants during the flight game and in the torso JAR with visual feedback suggest that these children attempt to resolve the visual discrepancy by compensatory head movements. This is likely due to the weaker reliability of the neck proprioception, which is not mature yet at this developmental stage (*54–56*), and which caused the visual inputs to be stronger weighted. This behavioral pattern aligns with recent works showing biases in the perception of visual and haptic verticality to unusual body orientations in younger children (*57, 58*), which is here confirmed by the younger participant’s inability to stabilize their head vertically while aligning their torso to lateral target positions. The stronger reliance on the visual system we observed in younger children has been shown to disappear in adults, where immersive VR appears to increase the contribution of proprioceptive and vestibular inputs to postural control over vision (*59*).

The joint display of these two behaviors led to the unexpected observation that only the older participants favorably selected an ‘en-bloc’ strategy with a stiff intersegmental link during the steering task. This opposes the accepted model of postural development, which states that such behavior is preferentially observed in younger children and decreases with age(*1, 13, 15*). One study found a similar behavior in adults, who displayed a head-to-torso stabilization in dimensions in which independent head movements were not beneficial (*14*). This is concomitant with our results, as head movements in the torso-controlled trials tended to disturb the participants’ spatial orientation. Younger children instead failed to use this simpler pattern, which suggests that the altered visual feedback provided by the VR setup interfered with the selection of an adequate coordination strategy, likely by reweighting the sensory contributions to posture estimation (*59*).

## Conclusion

We showed that the immersion in a virtual environment where the effects of head and torso are decoupled causes children aged 6 and 8 to deviate from an ‘en-block’ postural control, currently accepted as the default coordination strategy at these ages. This suggests that the still developing proprioception at the neck and torso levels and the strong reliance on visual feedback causes these children to overestimate their torso displacement and to correct the resulting visual discrepancy through compensatory head movements. We argue that, at this developmental stage, the postural control is not yet mature enough to be robust to an alteration of the visual input, which prevents an effective visual-vestibular-proprioceptive sensory integration, and confirms that the maturation of motor control extends beyond childhood.

## Methods

### Experimental Design

The objectives of the studies presented in this work were to (i): assess the ability of school-aged children to use and interact with an immersive virtual platform, steered by body movements, (ii) to compare this ability with the capacity displayed by healthy young adults, (iii) identify and describe the coordination patterns which emerge during the use of such a system, (iv) evaluate the development of these patterns along childhood, and (v) to disambiguate the contribution of the visual and proprioceptive systems to postural control and motor coordination during the use of the system described in (i).

The study was designed following a repeated measures design, where all participants were asked to use the platform using their head and their torso at multiple timepoints (see *Experimental Protocols* below for details), using the participants’ ages as a between-subjects factor. The participants were randomly assigned to start with their head or their torso. The sample sizes were determined using the software G*power(*60*), to reach a significance level of α = 0.05 and a power of (1-β) = 0.95.

### Subjects

Thirty-six typically developing children participated in the first study, grouped as follows: nine 6-year-olds (5 girls), eight 8-year-olds (2 girls), four 9-year-olds (1 girl) and eleven 10-year-olds (2 girls). Two children (aged 6 and 8) asked to stop the experiment and two other ones (aged 8 and 10) did not comply with the instructions; their data were excluded from further analyses. In addition, 13 healthy adults participated in the study (3 women, age 28.5±3.4 years). Twenty-four typically developing children participated in the second study, grouped as follows: ten 6-year-olds (7 girls), ten 8-year-olds (5 girls), and ten 10-year-olds (5 girls), as well as 10 healthy adults (4 women, age 27.0±3.2 years). Two 6-year-olds did not complete the session with the flight simulator, their data are reported only for the JAR task. Both studies were approved by the local ethical committees and were carried out in accordance with the Helsinki declaration. All the participants or their legal representative gave their written consent to take part in this study.

### Experimental setup

The participants were equipped with a head-mounted display (HMD, Oculus Rift) through which they were shown the virtual environment, and an inertial measurement unit (IMU, X-sens MTw Awinda) placed in their back between the scapulae and maintained with a custom harness to acquire their trunk’s 3-dimensional (3D) rotation (see Fig. 1B). The IMU embedded within the HMD was used both to control the view in the virtual environment and to acquire the head rotations. The kinematic data were acquired at a sample period of 68 ms.

### Virtual environment and navigation task

We created a virtual environment (VE) using the game engine Unity3D, which represented a FPV flight on a bird’s back at a constant speed of 12 m/s, (*44, 45*). A succession of coins to catch (distance between consecutive coins: 58m) represented a path to follow, randomly alternating simple forward motion and one of four directional maneuvers (right turn, left turn, ascent, descent). The coins’ initial diameter was 1 m, and every time one coin was caught, the next one was enlarged to 2 m. To minimize possible effects of path planning abilities, we additionally displayed a colored line smoothly connecting the coins, computed as a Catmull-Rom spline (*61*). Similarly, to provide the participants with a visual cue of their own position in space, an eagle was displayed below their visual horizon (see Fig. 1A). Finally, to keep the experiment engaging, a tinkling sound was played when the coin was caught at a distance smaller than 10 m, which also added points to a total score for the trial, displayed at the top of the screen.

### Control of the flight simulator

The participants were asked to control the flight simulator using either head or trunk movements. Ascent and descent were achieved by flexion and extension of the controlling body part while right and left turns were computed as a linear combination of lateral flexion and axial rotation. The head and torso rotations were reset to zero before each sequence, at the participants’ self-selected neutral position corresponding to a straight, forward flight. Continuous tracking of the head movements also enabled a dynamic adaptation of the field of view, allowing the users to look around in the virtual environment. Steering with torso movements, therefore, required decoupling vision and steering commands, whereas these aspects were tied in the head-controlled trials.

### Joint angle reproduction (JAR) task

We created a JAR task (*46–48*) in virtual reality using the game engine Unity 3D. The participants were immersed in a virtual landscape and were asked to align their head or their torso to one of three predefined orientations (0° and +/−15°) indicated by a pink line. We tested three conditions: *Feedback*, where a blue line showed the current orientation of the controlling body part, *Still*, where the additional visual feedback was removed and *Forward*, where a constant forward speed was simulated. The duration of one trial was set to 4 s, and the participants were asked to hold their final position until the next trial.

### Experimental protocol study 1

Upon arriving, the participants were shown the movements to control the simulator using the head or the torso. They were equipped with the HMD and the IMU, and were seated on a stool or on a chair and asked not to lean against the backrest. The participants were randomly allocated to start the experiment using the head or the torso, using adaptive covariate randomization with the gender as covariate (*62*). For the torso-controlled trials, the participants were advised to keep their neck rigid as to move their entire upper body as a whole. Similarly, before starting the head-controlled trials, the experimenter made the participants aware that moving their trunk was unnecessary.

The recording sessions took place on two consecutive days. On day 1, the participants had to steer the simulator along four paths with each body part. The first sequence contained 26 coins and was an initial evaluation of the performance (hereafter: *Before*). The second and third sequences each contained 50 coins; these sequences were considered as training. The fourth sequence contained 18 coins (hereafter: *After*). All the sequences controlled with a given body part were executed successively. On day 2, one sequence containing 26 coins had to be performed with each body part (hereafter: *Day After*). Breaks were allowed between the sequences, at the participants’ demand.

### Experimental protocol study 2

The participants were equipped and seated as previously and were shown the JAR movements by the experimenter. The conditions were tested in the following order: *Feedback, Still, Forward*, while the participants were randomly allocated to start either with the head or the torso, using covariate adaptive randomization with the gender as covariate (*62*). The orientations were presented in a randomized order, totalling 5 repetitions for each orientation in the *Feedback* condition and 10 repetitions for the *Still* and *Forward* conditions. At the end of the session, the participants executed one flight sequence with the simulator (*Before* session described above).

### Data processing

The kinematic data acquired in study 1 was divided into segments corresponding to the intervals between consecutive coins. Descriptive variables were computed on these segments and averaged over each entire sequence (see Table 1). Principal component analysis (PCA) was applied to the dataset containing the kinematic variables extracted from all trials, or from the head- and torso-controlled trials, respectively. Outliers were detected as data points whose Euclidean distance to the centroid of the z-scored dataset deviated from the average value by more than 4 standard deviations. These points were given a weight of 0.5 in the PCA computation. The variables with normalized loadings > 0.75 on the first (all trials, head-controlled trials) or the first two principal components (torso trials) were considered as significant and were regrouped into functional clusters.

**Table 1:** Descriptive kinematic variables.

| # | Variable | Details |
| --- | --- | --- |
| **Steering performance** |  |  |
| **1-3** | Error (a.u.) | Unsigned distance to the coin center, computed when the participant crossed the vertical plane perpendicular to the trajectory supporting the coin. |
| **4** | Path ratio (-) | Quotient of the travelled path and an ideal path computed as a Catmull-Rom interpolation between the coins. Computed for the entire sequence. |
| **5** | Time (s) | Duration of the interval between two consecutive coins. |
| **Head movements** |  |  |
| **6-8** | Head rotation amplitude (°) | Interquartile range. *Pitch, roll, yaw* |
| **Torso movements** |  |  |
| **9-11** | Torso rotation amplitude (°) | Interquartile range. *Pitch, roll, yaw* |
| **12-14,18** | Mean torso speed (°) | Angular velocity. *Pitch, roll, yaw, norm* |
| **15-17, 19** | Maximum torso speed (°) | Angular velocity. *Pitch, roll, yaw, norm* |
| **Head-torso coordination** |  |  |
| **20-24** | Head-torso correlation (-) | Absolute correlation.<br>*Pitch-pitch, roll-roll, yaw-yaw, roll-yaw, yaw-roll* |
| **25-27** | Head anchoring index (AI,-) | Computed as $\Delta\sigma = \frac{\sigma_r - \sigma_a}{\sigma_r + \sigma_a}$, where $\sigma_a$ is the standard deviation of the absolute head angles and $\sigma_r$ the standard deviation of the head angles relative to the torso. Positive $\Delta\sigma$ values indicate a preferred head stabilization to the external space and negative values a better head stabilization to the torso (9, 14). *Pitch, roll, yaw* |
| **28-32** | Peak time of head-torso cross-correlation (s) | Occurrence of the peak in cross-correlation. Negative delays indicate that the head is moving ahead of the body.<br>*Pitch-pitch, roll-roll, yaw-yaw, roll-yaw, yaw-roll* |
| **33-35** | DTW distance (a.u.) | Dynamic time warping (DTW) distance between the head and torso sequences. Both segments were linearly interpolated to keep the number of data points constant across sequences (64, 65).<br>*Pitch, roll, yaw* |
| **Movement smoothness** |  |  |
| **36-38** | Torso SAL | 3-dimensional smoothness metric based on the arc length of the movement speed profile's normalized Fourier magnitude spectrum; higher absolute values relate to jerkier movements(66) |
| **39-41** | Number of peaks head (-) | Time-normalized number of peaks (67). *Pitch, roll, yaw* |
| **42-44** | Number of peaks torso (-) | Time-normalized number of peaks (67). *Pitch, roll, yaw* |
| **45-47** | Number of peaks bird (-) | Time-normalized number of peaks (67). *Pitch, roll, yaw* |
| **48-50** | Torso speed ratio (-) | Ratio of the mean to the maximum velocities; a ratio close to 1 stands for smooth movements, while lower values indicate jerkier movements (68, 69). *Pitch, roll, yaw* |

The data acquired during Study 2 was separated into individual trials, and the final position was averaged over the last 1.5 s of each trial. For each trial, we computed the signed error with respect to the target orientation, the overshoot, the number of oscillations around the final angle, and for the trials involving the torso, the head AI (computed over the entire trial), the final angular difference of the head and the torso and the head alignment “error” as the difference between the final head angle and the target orientation.

### Statistical analysis

The statistical evaluations were performed using paired t-tests or repeated-measures ANOVAs, using the age as a between-subjects factor and the control type and/or experimental phase as within-subject factors using custom Matlab routines (*63*). The p-values were corrected using the Greenhouse-Geisser correction when Mauchly’s test indicated a violation of sphericity. Post hoc analyses were conducted using Tukey’s honest significant differences test, with a significance level of.05 for all tests.

## Acknowledgments

The authors thank Stefania Saviotti, Elisa Freddi and Davide Esposito for their help in the data collection.

## Author contributions

J.M., L.C., M.G., and S.M. designed the studies; J.M., L.G., and S.Z. enrolled the participants; J.M., L.C. and S.Z. performed the experiments; J.M. analyzed the data and wote the manuscript; J.M., L.C., M.G., and S.M. revised and edited the manuscript.

## Data and materials availability

All data needed to evaluate the conclusions in, the paper are present in the paper and/or the Supplementary Materials. Data used for this submission are available at https://zenodo.org/record/4314310#.X9IwzC2S2CM.

## Supplementary materials

**Table S1:** Post-hoc tests for the effect fn control type and experimental phase on the steering performance (error). Bold entries indicate comparisons with statistically significant differences at 0.05 level, * p < 0.05, ** p < 0.01, *** p <0.001

| Age | Control - Phase1 | Control - Phase2 | Difference | *p* | Cohen's *d* |
| --- | --- | --- | --- | --- | --- |
| 6 | Head - Before | Head - After | 6.1 | 0.113 | 0.6 |
|  | Head - Before | Head - Day After | 6.42 | 0.072 | 0.63 |
|  | **Head - Before** | **Torso - Before** | **-38.6** | **0.002** ** | **-1.17** |
|  | Head - After | Head - Day After | 0.32 | 0.693 | 0.12 |
|  | **Head - After** | **Torso - After** | **-14.27** | **0.01** ** | **-0.83** |
|  | **Head - Day After** | **Torso - Day After** | **-12.87** | **<0.001** *** | **-1.17** |
|  | **Torso - Before** | **Torso - After** | **30.43** | **0.013** * | **0.85** |
|  | **Torso - Before** | **Torso - Day After** | **32.15** | **0.014** * | **0.97** |
|  | Torso - After | Torso - Day After | 1.72 | 0.997 | 0.09 |
| 8 | Head - Before | Head - After | -0.07 | 1 | -0.14 |
|  | Head - Before | Head - Day After | 0.6 | 1 | 0.25 |
|  | **Head - Before** | **Torso - Before** | **-56.89** | **<0.001** *** | **-1.45** |
|  | Head - After | Head - Day After | 0.67 | 0.219 | 0.55 |
|  | Head - After | Torso - After | -6.3 | 0.83 | -3.8 |
|  | Head - Day After | Torso - Day After | -5.07 | 0.652 | -2.15 |
|  | **Torso - Before** | **Torso - After** | **50.51** | **0.001** ** | **1.26** |
|  | **Torso - Before** | **Torso - Day After** | **52.42** | **0.002** ** | **1.28** |
|  | Torso - After | Torso - Day After | 1.9 | 0.999 | 0.27 |
| 9 | Head - Before | Head - After | 1.48 | 0.998 | 1.04 |
|  | Head - Before | Head - Day After | 1.61 | 0.997 | 1.12 |
|  | Head - Before | Torso - Before | -7.75 | 0.992 | -1.25 |
|  | Head - After | Head - Day After | 0.13 | 0.998 | 0.28 |
|  | Head - After | Torso - After | -5.22 | 0.945 | -1.07 |
|  | Head - Day After | Torso - Day After | -3.51 | 0.932 | -1.34 |
|  | Torso - Before | Torso - After | 4.01 | 1 | 0.52 |
|  | Torso - Before | Torso - Day After | 5.85 | 0.998 | 0.89 |
|  | Torso - After | Torso - Day After | 1.84 | 0.999 | 0.33 |
| 10 | Head - Before | Head - After | 0.69 | 0.999 | 0.37 |
|  | Head - Before | Head - Day After | 1.21 | 0.991 | 0.7 |
|  | Head - Before | Torso - Before | -5.85 | 0.978 | -1.4 |
|  | Head - After | Head - Day After | 0.52 | 0.109 | 0.68 |
|  | Head - After | Torso - After | -6.3 | 0.485 | -1.08 |
|  | Head - Day After | Torso - Day After | -3.52 | 0.625 | -1.24 |
|  | Torso - Before | Torso - After | 0.24 | 1 | 0.03 |
|  | Torso - Before | Torso - Day After | 3.53 | 0.998 | 0.74 |
|  | Torso - After | Torso - Day After | 3.3 | 0.91 | 0.51 |
| Adults | Head - Before | Head - After | 0.42 | 1 | 1.25 |
|  | Head - Before | Head - Day After | 0.45 | 1 | 1.23 |
|  | Head - Before | Torso - Before | -0.63 | 1 | -0.67 |
|  | Head - After | Head - Day After | 0.03 | 1 | 0.11 |
|  | Head - After | Torso - After | -0.48 | 1 | -1.17 |
|  | Head - Day After | Torso - Day After | -0.54 | 1 | -0.6 |
|  | Torso - Before | Torso - After | 0.57 | 1 | 0.53 |
|  | Torso - Before | Torso - Day After | 0.54 | 1 | 0.39 |
|  | Torso - After | Torso - Day After | -0.03 | 1 | -0.02 |

**Table S2:** Post-hoc tests for the effect of age on the steering performance (error). Bold entries indicate comparisons with statistically significant differences at 0.05 level, * p < 0.05, ** p < 0.01, *** p <0.001

| Control - Phase | Age1 | Age2 | Difference | *p* | Cohen's *d* |
| --- | --- | --- | --- | --- | --- |
| Head - Before | 6 | 8 | 7.67 | 0.288 | 0.73 |
|  | 6 | 9 | 6.49 | 0.526 | 0.54 |
|  | 6 | 10 | 7.26 | 0.155 | 0.76 |
|  | 6 | Adults | 8.44 | 0.07 | 0.95 |
|  | 8 | 9 | -1.18 | 0.999 | -0.92 |
|  | 8 | 10 | -0.41 | 1 | -0.28 |
|  | 8 | Adults | 0.77 | 1 | 0.98 |
|  | 9 | 10 | 0.77 | 1 | 0.33 |
|  | 9 | Adults | 1.95 | 0.988 | 2.12 |
|  | 10 | Adults | 1.19 | 0.994 | 0.74 |
| Head - After | 6 | 8 | 1.5 | 0.37 | 0.75 |
|  | 6 | 9 | 1.87 | 0.235 | 0.78 |
|  | 6 | 10 | 1.84 | 0.06 | 0.91 |
|  | **6** | **Adults** | **2.76** | **0.002** ** | **1.55** |
|  | 8 | 9 | 0.37 | 0.996 | 0.6 |
|  | 8 | 10 | 0.34 | 0.992 | 0.28 |
|  | 8 | Adults | 1.26 | 0.51 | 3.41 |
|  | 9 | 10 | -0.02 | 1 | -0.03 |
|  | 9 | Adults | 0.9 | 0.832 | 2.9 |
|  | 10 | Adults | 0.92 | 0.587 | 1.29 |
| Head - Day After | 6 | 8 | 1.85 | 0.115 | 0.78 |
|  | 6 | 9 | 1.69 | 0.241 | 0.73 |
|  | **6** | **10** | **2.05** | **0.013** * | **1.13** |
|  | **6** | **Adults** | **2.47** | **0.002** ** | **1.44** |
|  | 8 | 9 | -0.17 | 1 | 0.09 |
|  | 8 | 10 | 0.19 | 0.999 | 0.76 |
|  | 8 | Adults | 0.62 | 0.908 | 1.67 |
|  | 9 | 10 | 0.36 | 0.99 | 0.84 |
|  | 9 | Adults | 0.79 | 0.847 | 2.07 |
|  | 10 | Adults | 0.43 | 0.942 | 1.15 |
| Torso - Before | 6 | 8 | -10.62 | 0.959 | -0.09 |
|  | 6 | 9 | 37.33 | 0.192 | 0.98 |
|  | **6** | **10** | **40.01** | **0.023** * | **1.34** |
|  | **6** | **Adults** | **46.41** | **0.006** ** | **1.66** |
|  | 8 | 9 | 47.95 | 0.099 | 1.02 |
|  | **8** | **10** | **50.63** | **0.015** * | **1.45** |
|  | **8** | **Adults** | **57.03** | **0.005** ** | **1.78** |
|  | 9 | 10 | 2.67 | 1 | 0.43 |
|  | 9 | Adults | 9.08 | 0.98 | 2.3 |
|  | 10 | Adults | 6.4 | 0.983 | 1.73 |
| Torso - After | 6 | 8 | 9.46 | 0.657 | 0.52 |
|  | 6 | 9 | 10.91 | 0.598 | 0.52 |
|  | 6 | 10 | 9.81 | 0.419 | 0.57 |
|  | **6** | **Adults** | **16.54** | **0.042** * | **1.09** |
|  | 8 | 9 | 1.45 | 1 | 0.33 |
|  | 8 | 10 | 0.35 | 1 | 0.05 |
|  | 8 | Adults | 7.08 | 0.83 | 4.99 |
|  | 9 | 10 | -1.1 | 1 | -0.14 |
|  | 9 | Adults | 5.63 | 0.937 | 1.81 |
|  | 10 | Adults | 6.73 | 0.714 | 1.21 |
| Torso – Day After | 6 | 8 | 9.65 | 0.197 | 0.73 |
|  | 6 | 9 | 11.04 | 0.152 | 0.84 |
|  | **6** | **10** | **11.39** | **0.02** * | **1.07** |
|  | **6** | **Adults** | **14.8** | **0.001** ** | **1.52** |
|  | 8 | 9 | 1.39 | 0.999 | 0.65 |
|  | 8 | 10 | 1.75 | 0.993 | 0.71 |
|  | 8 | Adults | 5.15 | 0.737 | 2.57 |
|  | 9 | 10 | 0.36 | 1 | 0.09 |
|  | 9 | Adults | 3.76 | 0.921 | 1.91 |
|  | 10 | Adults | 3.41 | 0.842 | 1.2 |

**Table S3:** ANOVA on variables selected after PCA on all trials. Bold entries indicate comparisons with statistically significant differences at 0.05 level, * p < 0.05, ** p < 0.01, *** p <0.001

| Effect | Variable | *p* | F | DOF1 | DOF2 | $\eta_p^2$ |
| --- | --- | --- | --- | --- | --- | --- |
| Control | **Torso Amplitude, roll** | **<0.001 \*\*\*** | **56.98** | **4** | **35** | **0.62** |
|  | **Torso Amplitude, pitch** | **<0.001 \*\*\*** | **128.91** | **4** | **35** | **0.79** |
|  | **Torso Amplitude, yaw** | **<0.001 \*\*\*** | **72.52** | **4** | **35** | **0.67** |
|  | **Mean Speed Torso, roll** | **<0.001 \*\*\*** | **73.84** | **4** | **35** | **0.68** |
|  | **Mean Speed Torso, pitch** | **<0.001 \*\*\*** | **119.41** | **4** | **35** | **0.77** |
|  | **Mean Speed Torso, yaw** | **<0.001 \*\*\*** | **97.61** | **4** | **35** | **0.74** |
|  | **Max Speed Torso, roll** | **<0.001 \*\*\*** | **76.45** | **4** | **35** | **0.69** |
|  | **Max Speed Torso, pitch** | **<0.001 \*\*\*** | **111.96** | **4** | **35** | **0.76** |
|  | **Max Speed Torso, yaw** | **<0.001 \*\*\*** | **85.77** | **4** | **35** | **0.71** |
|  | **Torso Speed norm, mean** | **<0.001 \*\*\*** | **131.52** | **4** | **35** | **0.79** |
|  | **Torso Speed norm, max** | **<0.001 \*\*\*** | **121.77** | **4** | **35** | **0.78** |
|  | **Head-torso correlation, roll-roll** | **<0.001 \*\*\*** | **283.23** | **4** | **35** | **0.89** |
|  | **Head-torso correlation, roll-yaw** | **<0.001 \*\*\*** | **1792.71** | **4** | **35** | **0.98** |
|  | **Head-torso correlation, yaw-roll** | **<0.001 \*\*\*** | **195.11** | **4** | **35** | **0.85** |
|  | **AI, roll** | **<0.001 \*\*\*** | **83.59** | **4** | **35** | **0.70** |
|  | **AI, pitch** | **<0.001 \*\*\*** | **196.70** | **4** | **35** | **0.85** |
|  | **Cross-correlation peak time, pitch-pitch** | **<0.001 \*\*\*** | **313.97** | **4** | **35** | **0.90** |
|  | **Cross-correlation peak time, roll-yaw** | **<0.001 \*\*\*** | **115.65** | **4** | **35** | **0.77** |
|  | **Cross-correlation peak time, yaw-roll** | **<0.001 \*\*\*** | **208.79** | **4** | **35** | **0.86** |
|  | **DTW distance, roll** | **<0.001 \*\*\*** | **231.23** | **4** | **35** | **0.87** |
|  | **DTW distance, yaw** | **<0.001 \*\*\*** | **161.63** | **4** | **35** | **0.82** |
| Age:Control | **Torso Amplitude, roll** | **0.017 \*** | **4.31** | **8** | **70** | **0.33** |
|  | Torso Amplitude, pitch | 0.057 | 3.12 | 8 | 70 | 0.26 |
|  | Torso Amplitude, yaw | 0.321 | 1.35 | 8 | 70 | 0.13 |
|  | Mean Speed Torso, roll | 0.086 | 2.67 | 8 | 70 | 0.23 |
|  | Mean Speed Torso, pitch | 0.073 | 2.81 | 8 | 70 | 0.24 |
|  | Mean Speed Torso, yaw | 0.342 | 1.29 | 8 | 70 | 0.13 |
|  | Max Speed Torso, roll | 0.228 | 1.67 | 8 | 70 | 0.16 |
|  | Max Speed Torso, pitch | 0.134 | 2.23 | 8 | 70 | 0.20 |
|  | Max Speed Torso, yaw | 0.240 | 1.62 | 8 | 70 | 0.16 |
|  | Torso Speed norm, mean | 0.208 | 1.79 | 8 | 70 | 0.17 |
|  | Torso Speed norm, max | 0.227 | 1.68 | 8 | 70 | 0.16 |
|  | **Head-torso correlation, roll-roll** | **0.002 \*\*** | **6.49** | **8** | **70** | **0.43** |
|  | Head-torso correlation, roll-yaw | 0.068 | 2.88 | 8 | 70 | 0.25 |
|  | **Head-torso correlation, yaw-roll** | **0.005 \*\*** | **5.37** | **8** | **70** | **0.38** |
|  | AI, roll | 0.068 | 2.92 | 8 | 70 | 0.25 |
|  | **AI, pitch** | **<0.001 \*\*\*** | **8.97** | **8** | **70** | **0.51** |
|  | **Cross-correlation peak time, pitch-pitch** | **0.036 \*** | **3.56** | **8** | **70** | **0.29** |
|  | Cross-correlation peak time, roll-yaw | 0.305 | 1.40 | 8 | 70 | 0.14 |
|  | **Cross-correlation peak time, yaw-roll** | **<0.001 \*\*\*** | **13.37** | **8** | **70** | **0.60** |
|  | **DTW distance, roll** | **<0.001 \*\*\*** | **9.66** | **8** | **70** | **0.52** |
|  | **DTW distance, yaw** | **<0.001 \*\*\*** | **9.24** | **8** | **70** | **0.51** |
| Phase:Control | Torso Amplitude, roll | 0.676 | 0.44 | 4 | 35 | 0.01 |
|  | Torso Amplitude, pitch | 0.145 | 2.49 | 4 | 35 | 0.07 |
|  | Torso Amplitude, yaw | 0.158 | 2.34 | 4 | 35 | 0.06 |
|  | Mean Speed Torso, roll | 0.514 | 0.79 | 4 | 35 | 0.02 |
|  | **Mean Speed Torso, pitch** | **0.012 \*** | **5.99** | **4** | **35** | **0.15** |
|  | **Mean Speed Torso, yaw** | **0.021 \*** | **5.29** | **4** | **35** | **0.13** |
|  | Max Speed Torso, roll | 0.374 | 1.13 | 4 | 35 | 0.03 |
|  | **Max Speed Torso, pitch** | **0.017 \*** | **5.94** | **4** | **35** | **0.15** |
|  | Max Speed Torso, yaw | 0.056 | 4.04 | 4 | 35 | 0.10 |
|  | **Torso Speed norm, mean** | **0.029 \*** | **4.75** | **4** | **35** | **0.12** |
|  | **Torso Speed norm, max** | **0.037** * | **4.72** | **4** | **35** | **0.12** |
|  | **Head-torso correlation, roll-roll** | **0.037** * | **4.38** | **4** | **35** | **0.11** |
|  | Head-torso correlation, roll-yaw | 0.060 | 3.97 | 4 | 35 | 0.10 |
|  | Head-torso correlation, yaw-roll | 0.086 | 3.16 | 4 | 35 | 0.08 |
|  | AI, roll | 0.132 | 2.67 | 4 | 35 | 0.07 |
|  | **AI, pitch** | **0.031** * | **4.68** | **4** | **35** | **0.12** |
|  | Cross-correlation peak time, pitch-pitch | 0.373 | 1.14 | 4 | 35 | 0.03 |
|  | Cross-correlation peak time, roll-yaw | 0.853 | 0.19 | 4 | 35 | 0.01 |
|  | Cross-correlation peak time, yaw-roll | 0.303 | 1.42 | 4 | 35 | 0.04 |
|  | DTW distance, roll | 0.143 | 2.44 | 4 | 35 | 0.07 |
|  | DTW distance, yaw | 0.137 | 2.59 | 4 | 35 | 0.07 |
| Age:Phase:Control | Torso Amplitude, roll | 0.531 | 0.95 | 8 | 70 | 0.10 |
|  | Torso Amplitude, pitch | 0.068 | 2.37 | 8 | 70 | 0.21 |
|  | Torso Amplitude, yaw | 0.218 | 1.56 | 8 | 70 | 0.15 |
|  | Mean Speed Torso, roll | 0.202 | 1.58 | 8 | 70 | 0.15 |
|  | **Mean Speed Torso, pitch** | **0.005** ** | **3.61** | **8** | **70** | **0.29** |
|  | Mean Speed Torso, yaw | 0.065 | 2.31 | 8 | 70 | 0.21 |
|  | Max Speed Torso, roll | 0.144 | 1.77 | 8 | 70 | 0.17 |
|  | Max Speed Torso, pitch | 0.129 | 1.94 | 8 | 70 | 0.18 |
|  | Max Speed Torso, yaw | 0.093 | 2.10 | 8 | 70 | 0.19 |
|  | **Torso Speed norm, mean** | **0.009** ** | **3.32** | **8** | **70** | **0.27** |
|  | Torso Speed norm, max | 0.105 | 2.07 | 8 | 70 | 0.19 |
|  | Head-torso correlation, roll-roll | 0.219 | 1.52 | 8 | 70 | 0.15 |
|  | Head-torso correlation, roll-yaw | 0.611 | 0.82 | 8 | 70 | 0.09 |
|  | Head-torso correlation, yaw-roll | 0.195 | 1.61 | 8 | 70 | 0.16 |
|  | AI, roll | 0.975 | 0.24 | 8 | 70 | 0.03 |
|  | AI, pitch | 0.253 | 1.42 | 8 | 70 | 0.14 |
|  | Cross-correlation peak time, pitch-pitch | 0.858 | 0.50 | 8 | 70 | 0.05 |
|  | Cross-correlation peak time, roll-yaw | 0.611 | 0.83 | 8 | 70 | 0.09 |
|  | Cross-correlation peak time, yaw-roll | 0.116 | 2.02 | 8 | 70 | 0.19 |
|  | DTW distance, roll | 0.533 | 0.94 | 8 | 70 | 0.10 |
|  | DTW distance, yaw | 0.925 | 0.38 | 8 | 70 | 0.04 |

**Table S4:** ANOVA on variables selected after PCA on the torso-controlled trials. Bold entries indicate comparisons with statistically significant differences at 0.05 level, * p < 0.05, ** p < 0.01, *** p <0.001

| Effect | Variable | *p* | F | DOF1 | DOF2 | $\eta_p^2$ |
| --- | --- | --- | --- | --- | --- | --- |
| Age | Error | **0.003** ** | **6.92** | **4** | **35** | **0.44** |
|  | Head Amplitude, pitch | **0.004** ** | **6.32** | **4** | **35** | **0.42** |
|  | Head Amplitude, roll | **0.004** ** | **5.92** | **4** | **35** | **0.4** |
|  | Torso Amplitude, roll | **0.041** * | **3.12** | **4** | **35** | **0.26** |
|  | Torso Amplitude, pitch | **0.003** ** | **6.94** | **4** | **35** | **0.44** |
|  | Torso Amplitude, yaw | **0.027** * | **3.67** | **4** | **35** | **0.3** |
|  | Mean Speed Torso, roll | 0.226 | 1.53 | 4 | 35 | 0.15 |
|  | Mean Speed Torso, pitch | **0.037** * | **3.31** | **4** | **35** | **0.27** |
|  | Mean Speed Torso, yaw | **0.041** * | **3.05** | **4** | **35** | **0.26** |
|  | Max Speed Torso, roll | 0.563 | 0.75 | 4 | 35 | 0.08 |
|  | Max Speed Torso, pitch | **0.037** * | **3.29** | **4** | **35** | **0.27** |
|  | Max Speed Torso, yaw | **0.016** * | **4.15** | **4** | **35** | **0.32** |
|  | Torso Speed norm, mean | 0.088 | 2.32 | 4 | 35 | 0.21 |
|  | Torso Speed norm, max | **0.048** * | **2.87** | **4** | **35** | **0.25** |
|  | Head-torso correlation, roll-roll | **0.01** * | **4.56** | **4** | **35** | **0.34** |
|  | Cross-correlation peak time, yaw-roll | **<0.001** *** | **9.57** | **4** | **35** | **0.52** |
|  | DTW distance, roll-roll | **0.005** ** | **5.7** | **4** | **35** | **0.39** |
|  | DTW distance, yaw-yaw | **0.003** ** | **6.89** | **4** | **35** | **0.44** |
| Age:Phase | Error | **0.007** ** | **4.51** | **8** | **70** | **0.29** |
|  | Head Amplitude, pitch | **0.007** ** | **3.45** | **8** | **70** | **0.1** |
|  | Head Amplitude, roll | **0.005** ** | **3.81** | **8** | **70** | **0.11** |
|  | Torso Amplitude, roll | **0.036** * | **2.62** | **8** | **70** | **0** |
|  | Torso Amplitude, pitch | **0.01** ** | **3.27** | **8** | **70** | **0.14** |
|  | Torso Amplitude, yaw | **0.041** * | **2.44** | **8** | **70** | **0.1** |
|  | Mean Speed torso, roll | **0.01** * | **3.27** | **8** | **70** | **0.02** |
|  | Mean Speed torso, pitch | **0.004** ** | **4** | **8** | **70** | **0.19** |
|  | Mean Speed torso, yaw | **0.009** ** | **3.36** | **8** | **70** | **0.21** |
|  | Max Speed torso, roll | **0.038** * | **2.51** | **8** | **70** | **0** |
|  | Max Speed torso, pitch | 0.087 | 1.93 | 8 | 70 | 0.14 |
|  | Max Speed torso, yaw | **0.045** * | **2.29** | **8** | **70** | **0.14** |
|  | Torso Speed norm, mean | **0.003** ** | **4.17** | **8** | **70** | **0.17** |
|  | Torso Speed norm, max | 0.085 | 1.97 | 8 | 70 | 0.11 |
|  | Head-torso correlation, roll-roll | 0.105 | 1.81 | 8 | 70 | 0.12 |
|  | Cross-correlation peak time, yaw-roll | **0.041** * | **2.38** | **8** | **70** | **0.07** |
|  | DTW distance, roll-roll | 0.088 | 1.92 | 8 | 70 | 0.14 |
|  | DTW distance, yaw-yaw | 0.263 | 1.3 | 8 | 70 | 0.11 |

**Table S5:** ANOVA on variables selected after PCA on the head-controlled trials. Bold entries indicate comparisons with statistically significant differences at 0.05 level, * p < 0.05, ** p < 0.01, *** p <0.001

| Effect | Variable | *p* | F | DOF1 | DOF2 | $\eta_p^2$ |
| --- | --- | --- | --- | --- | --- | --- |
| Age | Torso Amplitude, roll | 0.13 | 1.98 | 4 | 35 | 0.18 |
|  | **Torso Amplitude, pitch** | **0.016 \*** | **4.09** | **4** | **35** | **0.31** |
|  | **Torso Amplitude, yaw** | **0.015 \*** | **4.3** | **4** | **35** | **0.32** |
|  | Mean Speed torso, roll | 0.076 | 2.42 | 4 | 35 | 0.21 |
|  | **Mean Speed torso, pitch** | **0.015 \*** | **4.25** | **4** | **35** | **0.31** |
|  | **Mean Speed torso, yaw** | **0.015 \*** | **4.31** | **4** | **35** | **0.32** |
|  | Max Speed torso, roll | 0.068 | 2.55 | 4 | 35 | 0.22 |
|  | **Max Speed torso, pitch** | **0.015 \*** | **4.45** | **4** | **35** | **0.32** |
|  | **Max Speed torso, yaw** | **0.015 \*** | **4.81** | **4** | **35** | **0.34** |
|  | **Torso Speed norm, mean** | **0.016 \*** | **3.99** | **4** | **35** | **0.3** |
|  | **Torso Speed norm, max** | **0.015 \*** | **4.34** | **4** | **35** | **0.32** |
| Age:Phase | **Torso Amplitude, roll** | **0.018 \*** | **2.78** | **8** | **70** | **0.05** |
|  | **Torso Amplitude, pitch** | **0.038 \*** | **2.6** | **8** | **70** | **0.04** |
|  | **Torso Amplitude, yaw** | **0.016 \*** | **3.04** | **8** | **70** | **0.01** |
|  | **Mean Speed torso, roll** | **0.015 \*** | **3.21** | **8** | **70** | **0.11** |
|  | **Mean Speed torso, pitch** | **0.021 \*** | **2.92** | **8** | **70** | **0.07** |
|  | **Mean Speed torso, yaw** | **0.015 \*** | **3.26** | **8** | **70** | **0.11** |
|  | Max Speed torso, roll | 0.054 | 2.29 | 8 | 70 | 0.12 |
|  | Max Speed torso, pitch | 0.41 | 1.04 | 8 | 70 | 0.04 |
|  | **Max Speed torso, yaw** | **0.02 \*** | **2.7** | **8** | **70** | **0.1** |
|  | **Torso Speed norm, mean** | **0.015 \*** | **3.34** | **8** | **70** | **0.09** |
|  | Torso Speed norm, max | 0.316 | 1.23 | 8 | 70 | 0.05 |

**Table S6:** Post-hoc tests for the effect of control type and session on the JAR error. Bold entries indicate comparisons with statistically significant differences at 0.05 level, * p < 0.05, ** p < 0.01, *** p <0.001

| Age | Control – Session1 | Control – Session2 | Difference | *p* |  | Cohen's *d* |
| --- | --- | --- | --- | --- | --- | --- |
| 6 | Head - feedback | Head - still | 0.09 | 0.997 |  | 0.31 |
|  | Head - feedback | Head - forward | 0.31 | 0.632 |  | 0.82 |
|  | **Head - feedback** | **Torso - feedback** | **0.87** | **<0.001** | **\*\*\*** | **1.73** |
|  | Head - still | Head - forward | 0.22 | 0.525 |  | 0.52 |
|  | **Head - still** | **Torso - still** | **0.79** | **<0.001** | **\*\*\*** | **1.53** |
|  | **Head - forward** | **Torso - forward** | **0.41** | **0.017** | **\*** | **1.42** |
|  | Torso - feedback | Torso - still | 0.02 | 1.000 |  | -0.11 |
|  | Torso - feedback | Torso - forward | -0.14 | 0.920 |  | -0.34 |
|  | Torso - still | Torso - forward | -0.16 | 0.200 |  | -0.24 |
| 8 | Head - feedback | Head - still | -0.07 | 0.999 |  | -0.13 |
|  | Head - feedback | Head - forward | -0.05 | 1.000 |  | -0.10 |
|  | **Head - feedback** | **Torso - feedback** | **0.76** | **<0.001** | **\*\*\*** | **1.51** |
|  | Head - still | Head - forward | 0.02 | 1.000 |  | 0.04 |
|  | **Head - still** | **Torso - still** | **0.91** | **<0.001** | **\*\*\*** | **2.70** |
|  | **Head - forward** | **Torso - forward** | **0.88** | **<0.001** | **\*\*\*** | **2.72** |
|  | Torso - feedback | Torso - still | 0.08 | 0.982 |  | 0.26 |
|  | Torso - feedback | Torso - forward | 0.06 | 0.997 |  | 0.24 |
|  | Torso - still | Torso - forward | -0.02 | 1.000 |  | -0.07 |
| 10 | **Head - feedback** | **Head - still** | **0.77** | **0.004** | **\*\*** | **1.73** |
|  | **Head - feedback** | **Head - forward** | **0.78** | **0.003** | **\*\*** | **1.58** |
|  | **Head - feedback** | **Torso - feedback** | **1.01** | **<0.001** | **\*\*\*** | **1.91** |
|  | Head - still | Head - forward | 0.01 | 1.000 |  | 0.02 |
|  | **Head - still** | **Torso - still** | **0.66** | **<0.001** | **\*\*\*** | **2.97** |
|  | **Head - forward** | **Torso - forward** | **0.63** | **<0.001** | **\*\*\*** | **2.09** |
|  | **Torso - feedback** | **Torso - still** | **0.42** | **0.011** | **\*\*** | **1.18** |
|  | Torso - feedback | Torso - forward | 0.40 | 0.059 |  | 1.13 |
|  | Torso - still | Torso - forward | -0.02 | 0.999 |  | -0.18 |
| Adults | **Head - feedback** | **Head - still** | **1.47** | **<0.001** | **\*\*\*** | **2.84** |
|  | **Head - feedback** | **Head - forward** | **1.36** | **<0.001** | **\*\*\*** | **2.64** |
|  | **Head - feedback** | **Torso - feedback** | **1.58** | **<0.001** | **\*\*\*** | **3.24** |
|  | Head - still | Head - forward | -0.10 | 0.954 |  | -0.28 |
|  | **Head - still** | **Torso - still** | **0.84** | **<0.001** | **\*\*\*** | **3.11** |
|  | **Head - forward** | **Torso - forward** | **0.92** | **<0.001** | **\*\*\*** | **3.35** |
|  | **Torso - feedback** | **Torso - still** | **0.73** | **<0.001** | **\*\*\*** | **3.47** |
|  | **Torso - feedback** | **Torso - forward** | **0.71** | **<0.001** | **\*\*\*** | **3.33** |
|  | Torso - still | Torso - forward | -0.02 | 1.000 |  | -0.13 |

**Table S7:**
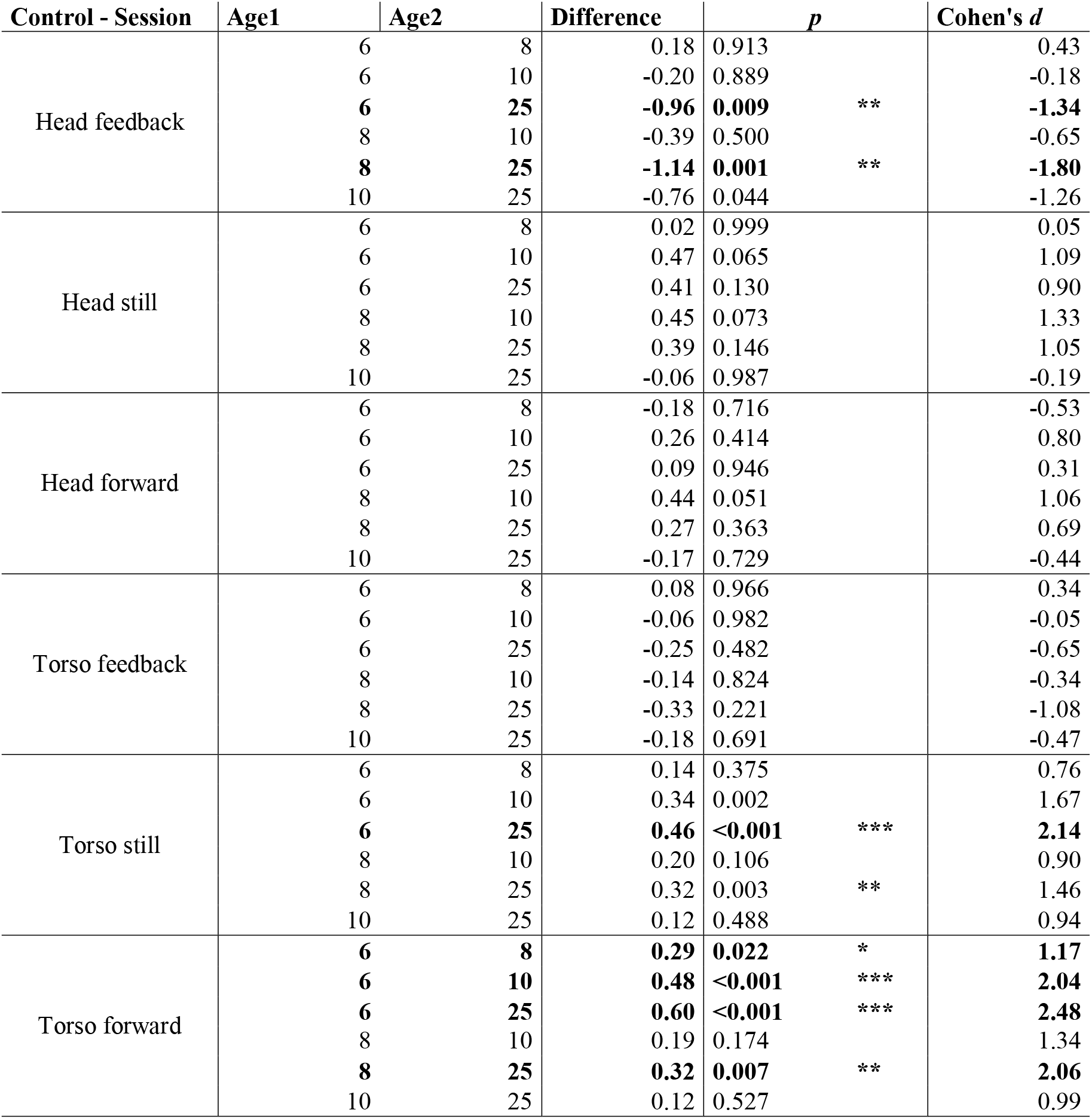
Post-hoc tests for the effect of age on the JAR error. Bold entries indicate comparisons with statistically significant differences at 0.05 level, * p < 0.05, ** p < 0.01, *** p <0.001

**Table S8:** Post-hoc tests for the effect of control type and session on the number of oscillations during the JAR test. Bold entries indicate comparisons with statistically significant differences at 0.05 level, * p < 0.05, ** p < 0.01, *** p <0.001

| Age | Control - Phase1 | Control - Phase2 | Difference | *p* |  | Cohen's *d* |
| --- | --- | --- | --- | --- | --- | --- |
| 6 | Head - feedback | Head - still | 0.09 | 0.997 |  | 0.31 |
|  | Head - feedback | Head - forward | 0.31 | 0.632 |  | 0.82 |
|  | **Head - feedback** | **Torso - feedback** | **0.87** | **<0.001** | **\*\*\*** | **1.73** |
|  | Head - still | Head - forward | 0.22 | 0.525 |  | 0.52 |
|  | **Head - still** | **Torso - still** | **0.79** | **<0.001** | **\*\*\*** | **1.53** |
|  | **Head - forward** | **Torso - forward** | **0.41** | **0.017** | **\*** | **1.42** |
|  | Torso - feedback | Torso - still | 0.02 | 1.000 |  | -0.11 |
|  | Torso - feedback | Torso - forward | -0.14 | 0.920 |  | -0.34 |
|  | Torso - still | Torso - forward | -0.16 | 0.200 |  | -0.24 |
| 8 | Head - feedback | Head - still | -0.07 | 0.999 |  | -0.13 |
|  | Head - feedback | Head - forward | -0.05 | 1.000 |  | -0.10 |
|  | **Head - feedback** | **Torso - feedback** | **0.76** | **<0.001** | **\*\*\*** | **1.51** |
|  | Head - still | Head - forward | 0.02 | 1.000 |  | 0.04 |
|  | **Head - still** | **Torso - still** | **0.91** | **<0.001** | **\*\*\*** | **2.70** |
|  | **Head - forward** | **Torso - forward** | **0.88** | **<0.001** | **\*\*\*** | **2.72** |
|  | Torso - feedback | Torso - still | 0.08 | 0.982 |  | 0.26 |
|  | Torso - feedback | Torso - forward | 0.06 | 0.997 |  | 0.24 |
|  | Torso - still | Torso - forward | -0.02 | 1.000 |  | -0.07 |
| 10 | **Head - feedback** | **Head - still** | **0.77** | **0.004** | **\*\*** | **1.73** |
|  | **Head - feedback** | **Head - forward** | **0.78** | **0.003** | **\*\*** | **1.58** |
|  | **Head - feedback** | **Torso - feedback** | **1.01** | **<0.001** | **\*\*\*** | **1.91** |
|  | Head - still | Head - forward | 0.01 | 1.000 |  | 0.02 |
|  | **Head - still** | **Torso - still** | **0.66** | **<0.001** | **\*\*\*** | **2.97** |
|  | **Head - forward** | **Torso - forward** | **0.63** | **<0.001** | **\*\*\*** | **2.09** |
|  | **Torso - feedback** | **Torso - still** | **0.42** | **0.011** | **\*** | **1.18** |
|  | Torso - feedback | Torso - forward | 0.40 | 0.059 |  | 1.13 |
|  | Torso - still | Torso - forward | -0.02 | 0.999 |  | -0.18 |
| Adults | **Head - feedback** | **Head - still** | **1.47** | **<0.001** | **\*\*\*** | **2.84** |
|  | **Head - feedback** | **Head - forward** | **1.36** | **<0.001** | **\*\*\*** | **2.64** |
|  | **Head - feedback** | **Torso - feedback** | **1.58** | **<0.001** | **\*\*\*** | **3.24** |
|  | Head - still | Head - forward | -0.10 | 0.954 |  | -0.28 |
|  | **Head - still** | **Torso - still** | **0.84** | **<0.001** | **\*\*\*** | **3.11** |
|  | **Head - forward** | **Torso - forward** | **0.92** | **<0.001** | **\*\*\*** | **3.35** |
|  | **Torso - feedback** | **Torso - still** | **0.73** | **<0.001** | **\*\*\*** | **3.47** |
|  | **Torso - feedback** | **Torso - forward** | **0.71** | **<0.001** | **\*\*\*** | **3.33** |
|  | Torso - still | Torso - forward | -0.02 | 1.000 |  | -0.13 |

**Table S9:** Post-hoc tests for the effect of age on the number of oscillations during the JAR test. Bold entries indicate comparisons with statistically significant differences at 0.05 level, * p < 0.05, ** p < 0.01, *** p <0.001

| Control - Phase1 | Age1 | Age2 | Difference | *p* |  | Cohen's *d* |
| --- | --- | --- | --- | --- | --- | --- |
| Head feedback | 6 | 8 | 0.18 | 0.913 |  | 0.43 |
|  | 6 | 10 | -0.20 | 0.889 |  | -0.18 |
|  | **6** | **25** | **-0.96** | **0.009** | **\*\*** | **-1.34** |
|  | 8 | 10 | -0.39 | 0.500 |  | -0.65 |
|  | **8** | **25** | **-1.14** | **0.001** | **\*\*** | **-1.80** |
|  | **10** | **25** | **-0.76** | **0.044** | **\*** | **-1.26** |
| Head still | 6 | 8 | 0.02 | 0.999 |  | 0.05 |
|  | 6 | 10 | 0.47 | 0.065 |  | 1.09 |
|  | 6 | 25 | 0.41 | 0.130 |  | 0.90 |
|  | 8 | 10 | 0.45 | 0.073 |  | 1.33 |
|  | 8 | 25 | 0.39 | 0.146 |  | 1.05 |
|  | 10 | 25 | -0.06 | 0.987 |  | -0.19 |
| Head forward | 6 | 8 | -0.18 | 0.716 |  | -0.53 |
|  | 6 | 10 | 0.26 | 0.414 |  | 0.80 |
|  | 6 | 25 | 0.09 | 0.946 |  | 0.31 |
|  | 8 | 10 | 0.44 | 0.051 |  | 1.06 |
|  | 8 | 25 | 0.27 | 0.363 |  | 0.69 |
|  | 10 | 25 | -0.17 | 0.729 |  | -0.44 |
| Torso feedback | 6 | 8 | 0.08 | 0.966 |  | 0.34 |
|  | 6 | 10 | -0.06 | 0.982 |  | -0.05 |
|  | 6 | 25 | -0.25 | 0.482 |  | -0.65 |
|  | 8 | 10 | -0.14 | 0.824 |  | -0.34 |
|  | 8 | 25 | -0.33 | 0.221 |  | -1.08 |
|  | 10 | 25 | -0.18 | 0.691 |  | -0.47 |
| Torso still | 6 | 8 | 0.14 | 0.375 |  | 0.76 |
|  | **6** | **10** | **0.34** | **0.002** | **\*\*** | **1.67** |
|  | **6** | **25** | **0.46** | **<0.001** | **\*\*\*** | **2.14** |
|  | 8 | 10 | 0.20 | 0.106 |  | 0.90 |
|  | **8** | **25** | **0.32** | **0.003** | **\*\*** | **1.46** |
|  | 10 | 25 | 0.12 | 0.488 |  | 0.94 |
| Torso forward | 6 | 8 | 0.29 | 0.022 |  | 1.17 |
|  | **6** | **10** | **0.48** | **<0.001** | **\*\*\*** | **2.04** |
|  | **6** | **25** | **0.60** | **<0.001** | **\*\*\*** | **2.48** |
|  | 8 | 10 | 0.19 | 0.174 |  | 1.34 |
|  | **8** | **25** | **0.32** | **0.007** | **\*\*** | **2.06** |
|  | 10 | 25 | 0.12 | 0.527 |  | 0.99 |

## References

1. C. Assaiante, S. Mallau, S. Viel, M. Jover, C. Schmitz, Development of Postural Control in Healthy Children: A Functional Approach. Neural Plast. 12, 109–118 (2005).

2. J. A. Ashton-Miller, K. M. McGlashen, A. B. Schultz, Trunk positioning accuracy in children 7-18 years old. J. Orthop. Res. 10, 217–225 (1992).

3. S. Mallau, M. Vaugoyeau, C. Assaiante, Postural Strategies and Sensory Integration: No Turning Point between Childhood and Adolescence. PLOS ONE. 5, e13078 (2010).

4. M. de Onis, WHO Motor Development Study: Windows of achievement for six gross motor development milestones. Acta Paediatr. 95, 86–95 (2006).

5. S. S. Yeo, S. H. Jang, S. M. Son, The different maturation of the corticospinal tract and corticoreticular pathway in normal brain development: diffusion tensor imaging study. Front. Hum. Neurosci. 8(2014), doi:10.3389/fnhum.2014.00573.

6. K. E. Adolph, J. M. Franchak, The development of motor behavior. Wiley Interdiscip. Rev. Cogn. Sci. 8(2017)., doi:10.1002/wcs.1430.

7. C. Simon-Martinez, G. L. dos Santos, E. Jaspers, R. Vanderschueren, L. Mailleux, K. Klingels, E. Ortibus, K. Desloovere, H. Feys, Age-related changes in upper limb motion during typical development. PLOS ONE. 13, e0198524 (2018).

8. J. C. van der Heide, B. Otten, L. A. van Eykern, M. Hadders-Algra, Development of postural adjustments during reaching in sitting children. Exp. Brain Res. 151, 32–45 (2003).

9. C. Assaiante, B. Amblard, Ontogenesis of head stabilization in space during locomotion in children: influence of visual cues. Exp. Brain Res. 93, 499–515 (1993).

10. C. Assaiante, B. Amblard, An ontogenetic model for the sensorimotor organization of balance control in humans. Hum. Mov. Sci. 14, 13–43 (1995).

11. N. A. Bernstein, The co-ordination and regulation of movements (Pergamon Press, Oxford, 1967).

12. O. Sporns, G. M. Edelman, Solving Bernstein’s Problem: A Proposal for the Development of Coordinated Movement by Selection. Child Dev. 64, 960–981 (1993).

13. M. N. Roncesvalles, C. Schmitz, M. Zedka, C. Assaiante, M. Woollacott, From egocentric to exocentric spatial orientation: development of posture control in bimanual and trunk inclination tasks. J. Mot. Behav. 37, 404–416 (2005).

14. H. Sveistrup, S. Schneiberg, P. A. McKinley, B. J. McFadyen, M. F. Levin, Head, arm and trunk coordination during reaching in children. Exp. Brain Res. 188, 237–247 (2008).

15. R. Grasso, C. Assaiante, P. Prévost, A. Berthoz, Development of Anticipatory Orienting Strategies During Locomotor Tasks in Children. Neurosci. Biobehav. Rev. 22, 533–539 (1998).

16. C. Chambers, T. Sokhey, D. Gaebler-Spira, K. P. Kording, The integration of probabilistic information during sensorimotor estimation is unimpaired in children with Cerebral Palsy. PLoS ONE. 12(2017)., doi:10.1371/journal.pone.0188741.

17. M. O. Ernst, M. S. Banks, Humans integrate visual and haptic information in a statistically optimal fashion. Nature. 415, 429–433 (2002).

18. C. Chambers, T. Sokhey, D. Gaebler-Spira, K. P. Kording, The development of Bayesian integration in sensorimotor estimation. J. Vis. 18(2018), doi:10.1167/18.12.8.

19. M. Gori, M. Del Viva, G. Sandini, D. C. Burr, Young Children Do Not Integrate Visual and Haptic Form Information. Curr. Biol. 18, 694–698 (2008).

20. D. Burr, M. Gori, in The Neural Bases of Multisensory Processes, M. M. Murray, M. T. Wallace, Eds. (CRC Press/Taylor & Francis, Boca Raton (FL), 2012), Frontiers in Neuroscience.

21. J. L. Contreras-Vidal, Development of forward models for hand localization and movement control in 6- to 10-year-old children. Hum. Mov. Sci. 25, 634–645 (2006).

22. J. Negen, B. Chere, L. Bird, E. Taylor, H. E. Roome, S. Keenaghan, L. Thaler, M. Nardini, Sensory Cue Combination in Children Under 10 Years of Age. bioRxiv, 501585 (2018).

23. M. Nardini, T. Dekker, K. Petrini, Crossmodal integration: a glimpse into the development of sensory remapping. Curr. Biol. CB. 24, R532–534 (2014).

24. S. Viel, M. Vaugoyeau, C. Assaiante, Adolescence: a transient period of proprioceptive neglect in sensory integration of postural control. Motor Control. 13, 25–42 (2009).

25. S. Greffou, A. Bertone, J.-M. Hanssens, J. Faubert, Development of visually driven postural reactivity: A fully immersive virtual reality study. J. Vis. 8, 15–15 (2008).

26. N. Gouleme, M. D. Ezane, S. Wiener-Vacher, M. P. Bucci, Spatial and temporal postural analysis: a developmental study in healthy children. Int. J. Dev. Neurosci. 38, 169–177 (2014).

27. P. J. Sparto, M. S. Redfern, J. G. Jasko, M. L. Casselbrant, E. M. Mandel, J. M. Furman, The influence of dynamic visual cues for postural control in children aged 7-12 years. Exp. Brain Res. 168, 505–516 (2006).

28. A. Shumway-Cook, M. H. Woollacott, The Growth of Stability. J. Mot. Behav. 17, 131–147 (1985).

29. M. Nardini, P. Jones, R. Bedford, O. Braddick, Development of Cue Integration in Human Navigation. Curr. Biol. 18, 689–693 (2008).

30. K. Petrini, A. Caradonna, C. Foster, N. Burgess, M. Nardini, How vision and self-motion combine or compete during path reproduction changes with age. Sci. Rep. 6, 29163 (2016).

31. A. Ramos, E. C. Hørning, I. L. Wilms, Simulated prism exposure in immersed virtual reality produces larger prismatic after-effects than standard prism exposure in healthy subjects. PLOS ONE. 14, e0217074 (2019).

32. J. M. Anglin, T. Sugiyama, S.-L. Liew, Visuomotor adaptation in head-mounted virtual reality versus conventional training. Sci. Rep. 7, 45469 (2017).

33. D. Cowie, A. McKenna, A. J. Bremner, J. E. Aspell, The development of bodily self-consciousness: changing responses to the Full Body Illusion in childhood. Dev. Sci. 21, e12557 (2018).

34. B. A. Morrongiello, M. Corbett, M. Milanovic, J. Beer, Using a Virtual Environment to Examine How Children Cross Streets: Advancing Our Understanding of How Injury Risk Arises. J. Pediatr. Psychol. 41, 265–275 (2016).

35. K. Y. Segovia, J. N. Bailenson, Virtually True: Children’s Acquisition of False Memories in Virtual Reality. Media Psychol. 12, 371–393 (2009).

36. E. Biffi, E. Beretta, A. Cesareo, C. Maghini, A. C. Turconi, G. Reni, S. Strazzer, An Immersive Virtual Reality Platform to Enhance Walking Ability of Children with Acquired Brain Injuries. Methods Inf. Med. 56, 119–126 (2017).

37. I. Bortone, D. Leonardis, N. Mastronicola, A. Crecchi, L. Bonfiglio, C. Procopio, M. Solazzi, A. Frisoli, Wearable Haptics and Immersive Virtual Reality Rehabilitation Training in Children With Neuromotor Impairments. IEEE Trans. Neural Syst. Rehabil. Eng. 26, 1469–1478 (2018).

38. C. B. de Mello Monteiro, T. Massetti, T. D. da Silva, J. van der Kamp, L. C. de Abreu, C. Leone, G. J. P. Savelsbergh, Transfer of motor learning from virtual to natural environments in individuals with cerebral palsy. Res. Dev. Disabil. 35, 2430–2437 (2014).

39. C. Gagliardi, A. C. Turconi, E. Biffi, C. Maghini, A. Marelli, A. Cesareo, E. Diella, D. Panzeri, Immersive Virtual Reality to Improve Walking Abilities in Cerebral Palsy: A Pilot Study. Ann. Biomed. Eng. 46, 1376–1384 (2018).

40. S. R. Sharar, G. J. Carrougher, D. Nakamura, H. G. Hoffman, D. K. Blough, D. R. Patterson, Factors Influencing the Efficacy of Virtual Reality Distraction Analgesia During Postburn Physical Therapy: Preliminary Results from 3 Ongoing Studies. Arch. Phys. Med. Rehabil. 88, S43–S49 (2007).

41. S. Won, J. Bailey, J. Bailenson, C. Tataru, I. A. Yoon, B. Golianu, Immersive Virtual Reality for Pediatric Pain. Children. 4(2017), doi:10.3390/children4070052.

42. H. Adams, G. Narasimham, J. Rieser, S. Creem-Regehr, J. Stefanucci, B. Bodenheimer, Locomotive Recalibration and Prism Adaptation of Children and Teens in Immersive Virtual Environments. IEEE Trans. Vis. Comput. Graph. 24, 1408–1417 (2018).

43. L. Tychsen, P. Foeller, Effects of Immersive Virtual Reality Headset Viewing on Young Children: Visuomotor Function, Postural Stability, and Motion Sickness. Am. J. Ophthalmol. 209, 151–159 (2020).

44. J. Miehlbradt, A. Cherpillod, S. Mintchev, M. Coscia, F. Artoni, D. Floreano, S. Micera, Data-driven body–machine interface for the accurate control of drones. Proc. Natl. Acad. Sci. 115, 7913–7918 (2018).

45. A. Cherpillod, D. Floreano, S. Mintchev, in 2019 Third IEEE International Conference on Robotic Computing (IRC) (2019), pp. 386–390.

46. S. Hillier, M. Immink, D. Thewlis, Assessing Proprioception: A Systematic Review of Possibilities. Neurorehabil. Neural Repair. 29, 933–949 (2015).

47. D. J. Goble, Proprioceptive Acuity Assessment Via Joint Position Matching: From Basic Science to General Practice. Phys. Ther. 90, 1176–1184 (2010).

48. D. J. Goble, C. A. Lewis, E. A. Hurvitz, S. H. Brown, Development of upper limb proprioceptive accuracy in children and adolescents. Hum. Mov. Sci. 24, 155–170 (2005).

49. H. Sigmundsson, H. T. A. Whiting, J. M. Loftesnes, Development of proprioceptive sensitivity. Exp. Brain Res. 135, 348–352 (2000).

50. S. Schneiberg, H. Sveistrup, B. McFadyen, P. McKinley, M. F. Levin, The development of coordination for reach-to-grasp movements in children. Exp. Brain Res. 146, 142–154 (2002).

51. L. H. C. Peeters, I. Kingma, G. S. Faber, J. H. van Dieën, I. J. M. de Groot, Trunk, head and pelvis interactions in healthy children when performing seated daily arm tasks. Exp. Brain Res. 236, 2023–2036 (2018).

52. T. Baumgartner, D. Speck, D. Wettstein, O. Masnari, G. Beeli, L. Jäncke, Feeling present in arousing virtual reality worlds: prefrontal brain regions differentially orchestrate presence experience in adults and children. Front. Hum. Neurosci. 2, 8 (2008).

53. L. Jäncke, M. Cheetham, T. Baumgartner, Virtual reality and the role of the prefrontal cortex in adults and children. Front. Neurosci. 3, 6 (2009).

54. T. Mergner, C. Siebold, G. Schweigart, W. Becker, Human perception of horizontal trunk and head rotation in space during vestibular and neck stimulation. Exp. Brain Res. 85, 389–404 (1991).

55. V. E. Pettorossi, M. Schieppati, Neck Proprioception Shapes Body Orientation and Perception of Motion. Front. Hum. Neurosci. 8, 895 (2014).

56. S. Hirabayashi, Y. Iwasaki, Developmental perspective of sensory organization on postural control. Brain Dev. 17, 111–113 (1995).

57. L. F. Cuturi, M. Gori, The Effect of Visual Experience on Perceived Haptic Verticality When Tilted in the Roll Plane. Front. Neurosci. 11, 687 (2017).

58. L. F. Cuturi, M. Gori, Biases in the Visual and Haptic Subjective Vertical Reveal the Role of Proprioceptive/Vestibular Priors in Child Development. Front. Neurol. 9(2019), doi:10.3389/fneur.2018.01151.

59. H. Akizuki, A. Uno, K. Arai, S. Morioka, S. Ohyama, S. Nishiike, K. Tamura, N. Takeda, Effects of immersion in virtual reality on postural control. Neurosci. Lett. 379, 23–26 (2005).

60. F. Faul, E. Erdfelder, A.-G. Lang, A. Buchner, G*Power 3: a flexible statistical power analysis program for the social, behavioral, and biomedical sciences. Behav. Res. Methods. 39, 175–191 (2007).

61. E. Catmull, R. Rom, in Computer Aided Geometric Design (New York: Academic Press, 1974), pp. 317–326.

62. J. W. Frane, A Method of Biased Coin Randomization, its Implementation, and its Validation. Drug Inf. J. 32, 423–432 (1998).

63. L. Caplette, Simple RM/Mixed ANOVA for any design. MATLAB Cent. File Exch., (available at https://www.mathworks.com/matlabcentral/fileexchange/64980-simple-rm-mixed-anova-for-any-design).

64. D. J. Berndt, J. Clifford, (AAAI, Seattle, WA, 1994; https://www.aaai.org/Papers/Workshops/1994/WS-94-03/WS94-03-031.pdf), vol. 10, pp. 359–370.

65. G. A. ten Holt, M. J. T. Reinders, E. A. Hendriks, (2007)., vol. 300.

66. S. Balasubramanian, A. Melendez-Calderon, E. Burdet, A Robust and Sensitive Metric for Quantifying Movement Smoothness. IEEE Trans. Biomed. Eng. 59, 2126–2136 (2012).

67. P. Gulde, J. Hermsdörfer, Smoothness Metrics in Complex Movement Tasks. Front. Neurol. 9(2018), doi:10.3389/fneur.2018.00615.

68. B. Rohrer, S. Fasoli, H. I. Krebs, R. Hughes, B. Volpe, W. R. Frontera, J. Stein, N. Hogan, Movement Smoothness Changes during Stroke Recovery. J. Neurosci. 22, 8297–8304 (2002).

69. L. Dipietro, H. I. Krebs, B. T. Volpe, J. Stein, C. Bever, S. T. Mernoff, S. E. Fasoli, N. Hogan, Learning, Not Adaptation, Characterizes Stroke Motor Recovery: Evidence From Kinematic Changes Induced by Robot-Assisted Therapy in Trained and Untrained Task in the Same Workspace. IEEE Trans. Neural Syst. Rehabil. Eng. 20, 48–57 (2012).

